# Microvilli regulate the release modes of alpha-tectorin to organize the domain-specific matrix architecture of the tectorial membrane

**DOI:** 10.1101/2024.01.04.574255

**Authors:** Ava Niazi, Ju Ang Kim, Dong-Kyu Kim, Di Lu, Igal Sterin, Joosang Park, Sungjin Park

## Abstract

The tectorial membrane (TM) is an apical extracellular matrix (ECM) in the cochlea essential for auditory transduction. The TM exhibits highly ordered domain-specific architecture. Alpha-tectorin/TECTA is a glycosylphosphatidylinositol (GPI)-anchored ECM protein essential for TM organization. Here, we identified that TECTA is released by distinct modes: proteolytic shedding by TMPRSS2 and GPI-anchor-dependent release from the microvillus tip. In the medial/limbal domain, proteolytically shed TECTA forms dense fibers. In the lateral/body domain produced by the supporting cells displaying dense microvilli, the proteolytic shedding restricts TECTA to the microvillus tip and compartmentalizes the collagen-binding site. The tip-localized TECTA, in turn, is released in a GPI-anchor-dependent manner to form collagen-crosslinking fibers, required for maintaining the spacing and parallel organization of collagen fibrils. Overall, we showed that distinct release modes of TECTA determine the domain-specific organization pattern, and the microvillus coordinates the release modes along its membrane to organize the higher-order ECM architecture.

Graphical Abstract

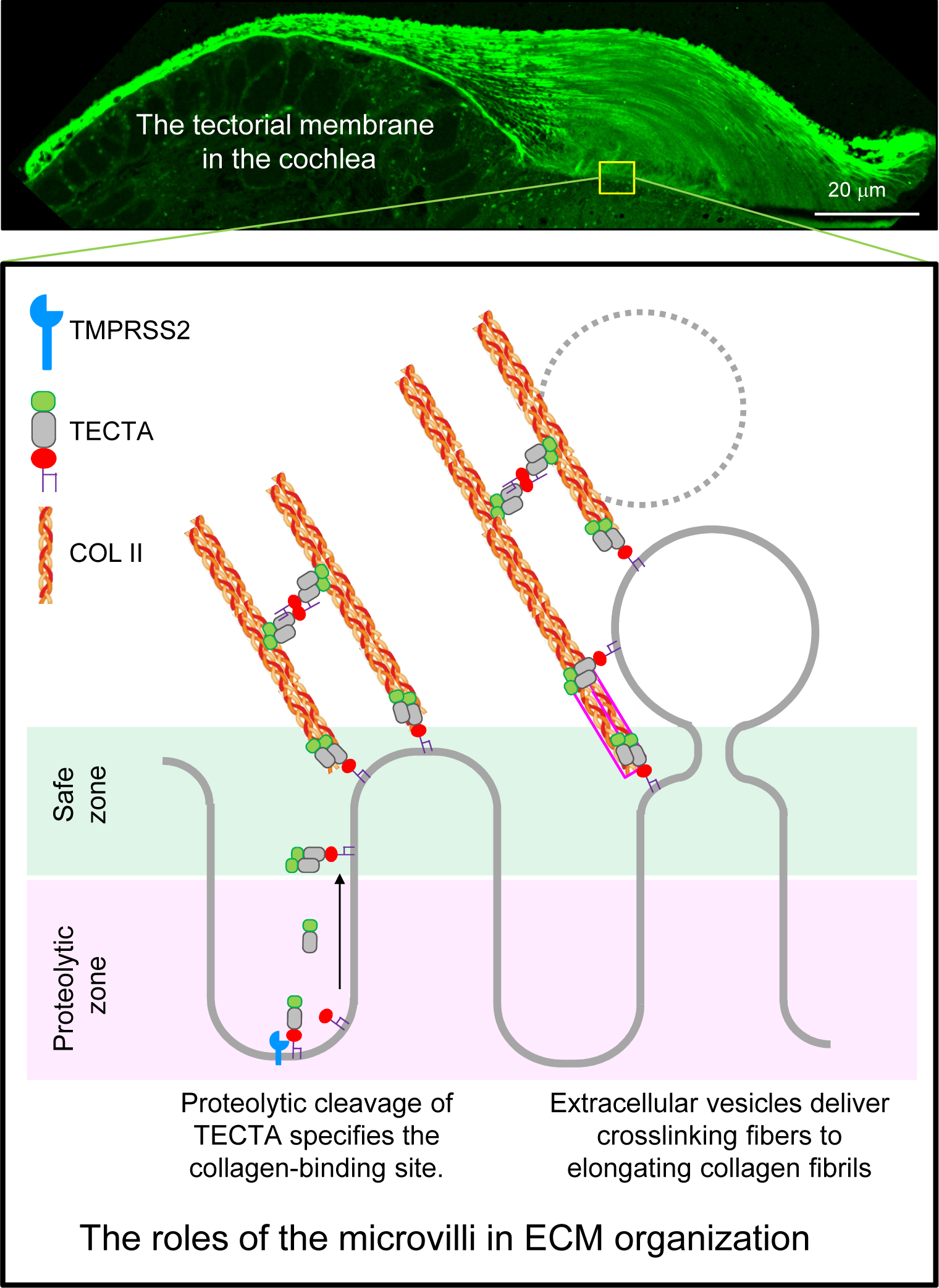

## Introduction

The extracellular matrix (ECM) maintains tissue architecture, transduces mechanical force, and regulates cellular signaling (Frantz et al.,2010). While many forms of ECMs exhibit highly ordered structural complexity (Peixoto and De Souza,1995; Richardson et al.,2007; Chen et al.,2015; Mouw et al.,2014; Holmes et al.,2018; Adams et al.,2023), how the specific ECM architecture is established outside the producing cells remains mysterious.

The tectorial membrane (TM) is an apical ECM generated by cochlear supporting cells and plays an essential role in coupling sound vibration to hair cell stereocilia (Goodyear and Richardson,2018). Abnormality of the TM causes sensorineural hearing loss (Legan et al.,2014; Ishai et al.,2019). The developing TM is organized on the apical membrane of cochlear supporting cells facing the lumen of the scala media and covers the entire organ of Corti (Rueda et al.,2003). The TM displays characteristic structural features: tonotopic, morphological, and stiffness gradient along the cochlear turn, structurally distinct domains along the radial axis, and differently oriented vertical layers (Andrade et al.,2016). α-tectorin/TECTA is a glycosylphosphatidylinositol (GPI)-anchored, zona pellucida (ZP) domain-containing protein and plays an essential role in TM organization (Goodyear and Richardson,2018); (Legan et al.,2000; Moreno-Pelayo et al.,2008; Kim et al.,2019). ZP domain-containing proteins are involved in the organization of the apical ECM in invertebrates and vertebrates, including the formation of the cuticle, the zona pellucida of the oocyte, and the luminal matrix fibers of the trachea and renal tubes (Jovine et al.,2005; Li Zheng et al.,2020; Wassarman and Litscher,2022; Drees et al.,2023; Stsiapanava et al.,2020). Unlike the basement ECM and interstitial matrix, which are organized within a confined space, the apical ECMs are assembled facing the luminal space or outside the body and often display structural complexity.

The medial part of the TM, called the limbal domain, is a dense matrix, which mediates the attachment of the TM to the spiral limbus. The lateral part of the TM, the body domain, displays an enlarged blade-like structure, which directly stimulates hair cell stereocilia. During development, collagen fibrils in the body domain are associated with the apical surface of the cochlear supporting cells. In the absence of surface-tethered TECTA, collagen fibrils are detached from the apical surface, which leads to severe disorganization of collagen fibrils (Kim *et al*., 2019). Surface-tethered TECTA is, in turn, released to compose the TM matrix. Notably, immuno-electron microscopy and high-resolution fluorescence imaging studies showed that released TECTA is organized into distinct shapes in each domain, forming dense fibers in the limbal domain and collagen-crosslinking fibers in the body domains, respectively (Andrade *et al*., 2016; Kim *et al*., 2019). While both surface-tethering and release of TECTA are essential for TM organization (Kim *et al*., 2019), how TECTA is specifically localized to the collagen-association site, how TECTA is released into the growing ECM, and how the released TECTA is organized into domain-specific shapes are unknown.

Notably, the shapes of the apical surface of cochlear supporting cells that produce each TM domain are strikingly different. The interdental cells of the spiral limbus, which produce the limbal domain, display a flat apical surface containing sparse microvilli, while that of the greater epithelial ridge (GER) cells, which produce the body domain matrix, display dense microvilli arrays, recruiting collagen fibrils on their tip membrane. Microvilli are actin-centered membrane protrusions on the apical surface of many types of epithelial cells (Sauvanet et al.,2015). The microvillus is a sorting station, and many proteins are differentially localized along its membrane (Jung et al.,2021; Ghosh et al.,2020). Interestingly, the matrix-association microdomain and membrane-adhesion microdomain are sharply segregated along the microvillus membrane (Sharkova et al.,2022; Houdusse and Titus,2021). For example, the pericellular matrix in the gut epithelial cells, glycocalyx, is associated with the microvillus tip, while the intermicrovillar adhesion complex composed of cell adhesion molecules is specifically localized on the lateral side (Crawley et al.,2014; Sun et al.,2020). In addition, microvilli are an extracellular vesicle (EV)-producing organelle, releasing EVs from the tip membrane (McConnell and Tyska,2007; McConnell et al.,2009; Choudhuri et al.,2014). EVs derived from the microvilli deliver signaling molecules, regulating tissue morphogenesis and immune response (Hurbain et al.,2022; Kim et al.,2018). However, how microvilli compartmentalize the matrix-association vs. membrane-adhesion domains and whether the budding of EVs from the microvillus tip regulates the organization of the associate matrix remain largely unclear.

In this study, we identified that TECTA is released by multiple mechanisms: proteolytic shedding and GPI-anchor-dependent release. Proteolytic shedding of TECTA is required to form dense fibers in the limbal domain. On the other hand, in the body domain, the proteolytic shedding of TECTA restricts surface-tethered TECTA to the microvillus tip, specifying the collagen-association site. The tip-localized TECTA is, in turn, released into the growing collagen fibrils in a GPI-anchor-dependent manner to form collagen-crosslinking fibers. The coordinated release of TECTA along the microvillus membrane plays a critical role in the domain-specific, higher-order matrix architecture.

## Materials and methods

### Animals and care

The animal care and procedures followed the standard regulations set by the National Institutes of Health and received approval from the Institutional Animal Care and Use Committee (Protocol 21-02004). The C57Bl6/J mice were kept in standard housing conditions at the University of Utah, with unrestricted access to food and water, and subjected to a 12:12 light/dark schedule.

### Cochlear tissue prep for semithin sections, Toluidine blue staining, TEM, and SEM

The cochlea was separated from the temporal bone following euthanasia (P2 or P28 mice). A minute aperture was created at the apex, and the sample was immersed in a fixative solution comprising 2.5% glutaraldehyde and 1% tannic acid in washing buffer [0.1 M sodium cacodylate (pH 7.4)] and left to incubate at 4°C overnight. Tissue samples collected at P28 were decalcified in 0.5 M EDTA (pH 8.0) for 7 days. After being rinsed, the cochlea underwent osmication (1% osmium tetroxide), dehydration (using ethanol and acetone), and saturation with Araldite/TAAB 812 resin kit (E202/1, TAAB Laboratories), in accordance with the manufacturer’s guidelines.

Briefly, the cochlea was sequentially immersed in a mixture of resin and acetone, starting with a 50:50% (v/v) ratio for 2 hours and then transitioning to 70:30% (v/v) overnight, before concluding with 100% resin incubation for an 8-hour to overnight duration. The plastic samples were then constructed within specimen embedding capsules (10504, Ted Pella) under a temperature of 60°C for 72 hours. Subsequently, these samples were sliced to a thickness of 1-μm using the Leica UC6 Ultramicrotome at the University of Utah’s Electron Microscopy Core Laboratory.

1. **Toluidine blue staining:** Each resulting section was briefly stained with a 100-μl droplet of 1% (w/v) Toluidine blue solution containing 1% (w/v) sodium borate and left on the heating block at the temperature of 60°C and allowed to air-dry prior to imaging.
2. **Transmission electron microscopy (TEM):** the plasticized tissues were cut further into radial sections of thickness ranging from 40 to 100 nm using a diamond knife on the Leica UC6 Ultramicrotome. The resulting sections were affixed onto copper grids (200 mesh) and subsequently stained in sequence with saturated aqueous uranyl acetate followed by Reynold’s Lead Citrate, for imaging.
3. **Scanning electron microscopy (SEM):** the fixed cochlear coil was gently separated from the bony shell and dehydrated serially. The samples were subjected to critical point drying (Pelco, CPD2) and mounted on a stub using a carbon sticky tab (Ted Pella, 16084-2). The samples were coated with gold/palladium using a sputter coater for imaging.

### Immunohistochemistry (IHC) of the cochlear tissue

The cochlea was separated from the temporal bone of the neonatal mice following euthanasia. A small opening was created at the apex of the cochlea, allowing fixative, a solution containing 4% paraformaldehyde (PFA) in phosphate-buffered saline (PBS), to access the inner compartments. The cochlea was then immersed in fixative at a temperature of 4°C overnight. Prior to OCT embedding, cryopreservation with sucrose was performed by placing the fixed cochlea in 15% sucrose in PBS until the tissue sinks, then transferring it to 30% sucrose in PBS until it sinks. Subsequently, the cochlea was immersed in a mixture of 50% optimal cutting temperature compound (OCT, 27050, Ted Pella) and 30% sucrose solution in PBS (v/v) at room temperature for 10 minutes, positioned in an embedding mold (18646A-1, Polysciences) containing 100% OCT, and frozen on dry ice. Thin sections of the frozen sample, approximately 7-μm thick, were cut and affixed to glass slides (22-037-246, Thermo Fisher Scientific). These tissue sections were then permeabilized in a PBST (0.2% Trition X-100) solution and blocked with a blocking buffer consisting of 2.5% normal goat serum and 2.5% normal donkey serum in PBST (containing 0.2% Triton X-100) for one hour at room temperature.

Antibody to TECTA was raised to peptides based on the predicted amino acid sequence of the mouse TECTA protein (CYNKNPLDDFLRPDGR) and was purified by antigen specific affinity purification by Abfrontier (www.gwvitek.com). The purified antibody (1 mg/ml) was used at working concentrations of 1:1000 dilution. Control experiments demonstrated that the antibody was, when used in isolation, specific for its respective antigens.

Following blocking, the tissue sections were incubated with primary antibodies including α-TECTA produced in rabbit (1:1000) or mouse α–Col II (1:100; ab150771, Abcam)) at 4°C overnight. Next, the tissues were washed with PBS and exposed to secondary antibodies including Alexa Fluor® 647 AffiniPure Donkey α-Rabbit IgG (H + L) (1:500), Cy3 AffiniPure Donkey Anti-Mouse IgG (H+L) (1:500), fluorescein isothiocyanate–PSA (at a concentration of 20 μg/ml; L0770, Sigma-Aldrich), Phalloidin-iFluor 594 Reagent (1:1000; ab176757, Abcam), and Hoechst 33342 (1:20,000) at room temperature for one hour. Subsequently, the tissues were washed with PBS three times, treated with ProLong™ Diamond Antifade Mountant (P36965, Thermo Fisher Scientific), and cover slipped for imaging.

Rabbit α–TMPRSS2 antibody (1:200; ab214462, Abcam) and mouse α–Annexin V antibody (1:250; sc-74438, Santa Cruz Biotechnology) required antigen retrieval. Tissues were incubated in citrate buffer pH 6 at 110 °C for 15 min, followed by blocking and antibody application as described above.

### RNAscope *in situ* hybridization (ISH)

Cryosectioned tissue samples were prepared as described above and RNAscope was performed by RNAscope® H202 & Protease Plus Reagents (#322330, ACDbio), RNAscope® 2.5 HD Duplex Detection Reagents (#322500, ACDbio) according to the manufacturer instruction. For *Tmprss1/Hpn* and *Tmprss2* detection RNAscope® Probe - Mm-Hpn (#852642, ACDbio) and RNAscope® Probe-Mm-Tmprss2 (#496721, ACDbio) were used respectively.

### Plasmid constructs

The pBK plasmid expressing a mouse form of TECTA (NM_001324548.1) was generously provided by Dr. Guy P. Richardson at the University of Sussex, UK, where single MYC (EQKLISEEDL) or HA (YPYDVPDYA) epitope is inserted at the N-terminal following the TECTA signal peptide (SP). Other plasmids were generated by PCR-based cloning.

1. **pBK/TECTA-ZP**: The C-terminus ZP domain of the TECTA (TECTA-ZP) was cloned using PCR with a primer set of 5’-AAACCGCGGGACCCTTGTGTGGGGGCG-3’ (forward primer) and 5’-AAACTCGAGTTATGAGGTTGCGCC-3’ (reverse primer). The PCR amplicon was cloned into pBK vector containing the TECTA SP and an epitope tag using SacII and XhoI.
2. **pBK/TECTA^R2061S^**: To genertat pBK/TECTA^R2061S^ a missense mutation was introduced in CCS1 (S<u>R</u>↓IA to S<u>S</u>↓IA: R2061S) site using overlap PCR with two primers sets of 5’-AAAGATATCAAGATCTCCTTAGACTCTG-3’ (forward primer of set 1), 5’-ATAATC TGTGGCGATGCTGGAATTTTGTGG-3’ (reverse primer of set 1), 5’-CCACAAAATTCCAGCATCGCCACAGATTAT-3’ (forward primer of set 2), and 5’-AAACTCGAGTTATGAGGTTGCGCC-3’ (reverse primer of set2). The second PCR amplicon was subcloned into the pBK/TECTA vector using EcoRV and XhoI restriction enzymes.
3. **pBK/TECTA-ZP^ΔGPI^ and pBK/TECTA-ZP^Clv^**: To generate pBK/TECTA-ZP^ΔGPI^ forward primer of set 1 and 5’-CTCGAGTCAGCTCCTTCCATCTTCTTGCAG-3’ (dGPI-R) were PCR. Similarly, and for pBK/TECTA-ZP^Clv^, forward primer of set 1 and 5’-AAACTCGAGTCACCTGGAATTTTGTGGGCA-3’ were used. PCR amplicons were subcloned into the pBK/TECTA vector using EcoRV and XhoI restriction enzymes.
4. **pBK/TECTA-ZP^ZP3^**: For generating pBK/TECTA-ZP^ZP3^, cDNA encoding ZP3 was restored from the mouse ovarian tissue by RT-PCR using PrimeScript RT Reagent Kit (Takara, RR037). Two PCR amplicons including the 5′ region of the TECTA-ZP and the transmembrane domain (TMD) of ZP3 were amplified primer pairs, 5′-CCGCGGGACCCTTGTGTGGGGGCG-3′ (nHA-WT-ZP-F), 5′-CTGCAAGATGGAAGGAGCTGGACTGCTTCT-3′ and 5′-CTGCAAGAAGATGGAAGGAGCTGGACTGCTT CT-3′ (mZP-Tectasec-F), 5′-CTTGTATCCCTTCCGCAATAACTCGA-3′ (XhoI-ZP3-R) respectively. An overlap extension PCR was performed using the amplicons and primers nHA-WT-ZP-F and XhoI-ZP3-R to produce chimeric DNA, which was further sub-cloned into pBK/TECTA using and EcoRV and XhoI sites.
5. **pCAGGS/TMPRSS2, pCAGGS/TMPRSS5, pCAGGS/PRSS36, pCAGGS/CORIN, pCAGGS/CTSK, and pCAGGS/HEPSIN:** Cochlea tissues were dissected from pooled wildtype mice (n=2) at P0. Total RNA was extracted using TRI Reagent (Invitrogen, AM9738). 500 ng of RNA template was subjected to in vitro reverse transcription using PrimeScript RT Reagent Kit (Takara, RR037). After elution with 40 ul of DNase/RNAse free water, 1 ul of the cDNA was used for PCR reaction using CloneAmp HiFi PCR Premix (Takara, 639298) with the following primers: 5’-CGGCTCGAGACCATGGCATTGAACTCAGGGTCF-3’ and 5’-CGGGGCGCGCCTTAGCTGTTCGCCCTCATTTGCT-3’ for TMPRSS2 (NM_015775); 5’-GCCAAGCTTACCATGAGTCCAACACTGGATGACCAAAGCCCAATGGAGATTCGGTGCACGGAAG-3’ and 5’-AATGGCGCGCCCTAGCGGACCTGCACAGTGTCATG-3’ for TMPRSS5 (NM_001359460); 5’-CGCGGCCGGCCACCATGTCCCATCACCTGTTTCTC-3’ and 5’-GGTCGACTGTAGTCTCAGCGAGGGATCAGTGGAGC-3’ for PRSS36 (NM_001319147); 5’-CGCGGCCGGCCACCATGGGCAGGGTTTCCTTCAG-3’ and 5’-CCGGGCGCGCCTTATCCTTGGGATTTCTTTTGGAG-3’ for CORIN (NM_016869); 5’-CGCGGCCGGCCACCATGTGGGTGTTCAAGTTTCTGCTG-3’ and 5’-CCGGGCGCGCCTCACATCTTGGGGAAGCTG-3’ for CTSK (NM_007802); 5’-AAAAAGCTTACCATGGCGAAGGAGGATGAGG-3’ (mHepsin-F) and 5’-AAAGGCGCGCCCTACTTGTCGTCGTCGTCCTTGTAGTCTTG GGGCTGAGTCACCATGCCACT-3’ for HEPSIN (NM_001110252.2). The PCR amplicons were subcloned into pCAGGS-null vector after restriction enzyme digestion at the sites of XhoI/AscI for TMPRSS2, HindIII/AscI for TMPRSS5 and HEPSIN, FseI/PshAI for PRSS36, and FseI/AscI for CORIN and CTSK using T4 ligase (NEB, M0202).

All correct clones were isolated by Sanger sequencing. All restriction enzymes were purchased from New England Biolabs and oligonucleotides used in this study were synthesized by the DNA/Peptide Facility, at the Health Sciences Center Cores at the University of Utah. BMP4 and Furin constructs were generously provided by Dr. Jan Christian at the university of Utah (Tilak et al.,2014).

### Cell culture and release assay

HEK293T cells were cultured in DMEM (Invitrogen, 11965092) containing 10% FBS (Invitrogen, 16140071), 1% Penicillin-Streptomycin (Invitrogen, 15140-122), and 1 mM sodium pyruvate (Invitrogen, 11360070). The cells were plated into 12-well plates (0.08 x 10^6^ cells/well) and transfected with each mTECTA or BMP4 plasmids (1.5 μg/well) and releasing enzymes including 0.12 μg/well of mGDE3, mGDE3H230A, mTMPRSS2, and mTMPRSS2S441A, Furin or 0.03 μg/well of HEPSIN. After 24 hours, culture media was replaced with fresh OptiMEM, and cells were further incubated for 16 hours. 1 μl PI-PLC derived from B. cereus (Thermo Fisher Scientific, P6466, 100 U/ml) was applied to culture media for 1 hour and media was then collected to measure TECTA release. The cells were then lysed with Cytosolic fraction buffer A (10 mM HEPES, 1.5 mM MgCl_2_, 10 mM KCL, 0.5 mM DTT, 0.05 % NP40, and protease inhibitors) at 4 °C for 30 min. Lysates were collected and spun down at 13,000 rpm at 4 °C for 30 min. The supernatant was then transferred to a new e-tube. Protein samples collected from culture media were treated with *PNGase F* enzyme (New England Biolabs, P0709S) to remove the *N*-linked oligosaccharides from the TECTA protein according to the manufacturer protocol. All protein samples were treated with Gel Loading Buffer (1xGLB, 6% Glycerol (v/v), 2% Sodium dodecyl sulfate (w/v), 1% Bromophenol Blue, 5% 2-Mercaptoethanol, 50 mM Tris-HCl, pH 6.8) before running SDS-PAGE.

### SDS-PAGE and western blot

Protein samples either from cells were loaded in SDS-PAGE gel in electrophoresis buffer (2.5 mM Tris-base, 19.2 mM Glycine and 0.1% SDS) at 120 V for 2 h. Proteins were transferred to the PVDF membrane in transfer buffer (2.5 mM Tris-base, 19.2 mM Glycine and 20% MeOH) at 100 V for 1 hour 30 min. The membrane was blocked in TBST (100 mM Tris-HCl, 150 mM NaCl, 0.1% Tween-20, pH 7.5) containing 5% skim milk for 1 hour and incubated at 4 °C overnight with antibodies including α-c-Myc produced in rabbit (Sigma-Aldrich, C3956, 1:2,000), and α-HA high affinity produced in rat (Sigma-Aldrich, 11867423001, 1:1,000). After washing with TBST, the membranes were incubated with a peroxidase-AffiniPure Donkey α-Rabbit IgG (H+L) (Jackson ImmunoResearch, 711-035-152, 1:5,000) or Peroxidase AffiniPure Goat Anti-Rat IgG (H+L) (Jackson ImmunoResearch, 112-035-003, 1:10,000) at RT for 1 hour. For exposure, the membrane was rinsed with TBST and was subjected to SuperSignal TM West Pico PLUS Chemiluminescent Substrate (Thermo Fisher Scientific, PI34580) or SuperSignalTM West Femto Chemiluminescent Substrate (Thermo Fisher Scientific, PI34095) and analyzed on a ChemiDocTM XRS+ system (Bio-Rad).

### Co-immunoprecipitation assay

HEK293T cells were cultured and plated (0.3 x 106 cells/well) in a 6-well plate that was coated with 25 μg/ml polyethyleneimine (PEI) for 1 hour prior to the cell seeding. The cells were transfected with 1.5 μg of Myc- and HA-mTECTA and 0.24 μg of GDE3 or TMPRSS2 constructs for 24 hrs. Cultured media was changed with 1 mL of OptiMEM for each well and further incubated for 24 hrs. Conditioned media was collected and centrifuged to remove pellet, and 1x protease inhibitor was added. 40 μl of supernatant was saved for input sample and 5 μl was used after treating 1xGLB for SDS-PAGE. Supernatant was transferred to a new micro-tube and precleared using 10 μl of protein A/G bead (Santa Cruz Biotechnology, sc-2003). It was then incubated with 0.5 μg of anti-Myc antibody for 1 hr at RT and incubated 20 μl of protein A bead for O/N at 4 °C. Immuno-precipitated beads were washed with 0.2% Triton X-100 in PBS for twice. 40 μl of 2xGLB without 2-Mercaptoethanol was used for elution. Eluent was separated from beads using centrifugation (13000 rpm, 5 min, RT) and 5% of 2-Mercaptoethanol was added. 15 μl of each eluent was used for SDS-PAGE.

### Surface biotinylation

HEK293T cells were cultured and plated (0.3 x 10^6^ cells/well) in a 6-well plate that was coated with 25 μg/ml polyethyleneimine (PEI) for 1 hour prior to the cell seeding. 4 μg each expression plasmid was transfected to each well in 500 μl of OptiMEM (Invitrogen, 31985070) and 10 μl Lipofectamine™ 2000 Transfection Reagent (Thermo Fisher Scientific, 11668019). After 24 hours, cells were washed with PBS and media was replaced with fresh OptiMEM and incubated for 16 hours. 1.5 μl PI-PLC was treated for 1 hour before harvest. The culture media were collected for the measurement of PBK/TECTA^ZP3^ release and cells were subjected to surface biotinylation. In brief, cells were washed with ice-cold PBS++ (1 mM CaCl2, 0.5 mM MgCl2 in PBS, pH 7.4) and incubated with 1 mg/ml EZ-LinkTMSulfo-NHS-SS-Bioctin solution (Thermo Fisher Scientific, 21331) at 4 °C for 40 min. Cells were washed with ice-cold PBS++ and the unreacted biotin was quenched with ice-cold glycine (100 mM) solution. Cells were then rinsed with ice-cold PBS++ and lysed with RIPA buffer (0.5% sodium deoxycholate, 0.1% sodium dodecyl sulfate, 1% TritonTM X-100, and protease inhibitors in PBS) at 4 °C for 20 min. Lysates were then sonicated and spin down with 13,000 rpm at 4 °C for 20 min. The supernatant was transferred to the new e-tube and collected for the “Total” fraction. To facilitate avidin binding, the supernatant was mixed with NeutrAvidinTM Agarose (Thermo Fisher Scientific, 29200) in a 1:1 ratio (v/v) within RIPA buffer. This mixture was then gently rotated at 4°C for 2 hours.

Subsequently, the Avidin-bound biotinylated protein were separated through centrifugation at 8000 rpm for 1 minute. Following two washes with RIPA buffe, the resulting material was collected as the “Biotin” fraction. All protein samples were treated with Gel Loading Buffer (GLB, 25% Glycerol (v/v), 2% Sodium dodecyl sulfate (w/v), 0.5 mg/ml Bromophenol Blue, 5% 2-Mercaptoethanol, 60 mM Tris-HCl, pH 6.8) before running SDS-PAGE.

### Live cell surface labeling

LS174T-W4 cells are human colon-derived epithelial cells that have been stably transfected with LKB1 and tetracycline-controlled STRAD, an LKB1-specific adaptor protein, expression vectors. This design allows for the activation of LKB1 by inducing STRAD expression through the addition of doxycycline (Hartmann et al.,2022) (Baas et al.,2004). These cells were generously provided by Dr. Willem-Jan Pannekoek (Center for Molecular Medicine, Utrecht, The Netherlands). The cells were cultured in RPMI 1640 medium (Invitrogen, 21870076) containing 10% tetracycline-free fetal calf serum (FCS) (TaKaRa Bio, 631106), 2 mM L-Glutamine (Thermo Fisher Scientific, 25030081), and 1% Penicillin-Streptomycin (Invitrogen, 15140-122). Activation of LKB1 and the corresponding expression of STRAD were induced by the addition of doxycycline (Sigma, D9891-1G) at a concentration of 1 μg/ml for 16 hours.

Cells were plated onto 24-well plate (0.08 x 10^6^ cells/well) containing 12 mm cover glass (Carolina Biological, 63<u>3009</u>) and were transfected with 0.8 μg of each pBK/TECAT-ZP^WT^, pBK/TECTA-ZP^R2061S^, and pCAGGS/TMPRSS2 with 50 μl OptiMEM, and 1.5 μl Lipofectamine™ 2000 Transfection Reagent Transfection Reagent. The next day after transfection, culture media was replaced with serum-free RPMI 1640 medium mixed 1 μg/ml doxycycline to allow induction of microvilli and the cells were left to grow for 16 hours. The cells were then washed and blocked for 1 min with a blocking buffer containing serum-free RPMI 1640, 10 mM HEPES (Thermo Fisher Scientific, 15630080), 2.5 % goat serum and 2.5 % donkey serum at RT. Following blocking, the cells were incubated with the primary antibody intended for surface staining, α-HA antibody produced in rat (MilliporeSigma, 11867423001, 1:150), at RT for 15 min. The cells were washed for 1 min with a RPMI 1640 solution containing, 10 mM HEPES three times and fixed with fixed with 4% PFA and 4% sucrose in PBS for 20 min. After quick wash with PBS, the cells were permeabilized with 0.2% Triton X-100 in PBS for 10 min and further blocked with a blocking solution (2.5 % normal goat serum, 2.5 % donkey serum in PBST containing 0.2% Triton X-100) and incubated with α-EPS8 antibody produced in mouse (BD Transduction Laboratories™, 610143, 1:50) at 4 °C overnight. After washing, cells were treated with appropriate secondary antibodies including Alexa Fluor® 488 AffiniPure Donkey Anti-Mouse IgG (H+L) (Jackson Immuno Research, 715-545-151, 1:1,000), Cy™3 AffiniPure Donkey Anti-Rat IgG (H+L) (Jackson Immuno Research, 711-165-152, 1:500), Phalloidin-iFluor 594 Reagent (1:1000; ab176757, Abcam), and Hoechst 33342 (1:20,000) for one hour at RT. Cells were then washed with PBS and mounted on the slide glass using ProLong™ Diamond Antifade Mountant (P36965, Thermo Fisher Scientific) in preparation for imaging.

### HEK-293 cell transfer assay

HEK293T cells were cultured and plated (0.3 x 10^6^ cells/well) in a 24-well plate that was coated with 25 μg/ml polyethyleneimine (PEI) for 1 hour prior to cell seeding. The cells were separated into two groups of donor and receiving cells. 0.8 μg each PBK/TECTA^WT^ and PBK/TECTA^RS^ expression plasmids and 0.1 μg PCAGGS/TMPRSS2 plasmid were transfected to each well using 400 μl OptiMEM (Invitrogen, 31985070) and 2 μl Lipofectamine™ 2000 Transfection Reagent (Thermo Fisher Scientific, 11668019). After 24 hours, cells were washed with PBS and media was replaced with fresh OptiMEM and incubated for 16 hours. 1.5 μl PI-PLC derived from B. cereus (Thermo Fisher Scientific, P6466, 100 U/ml) was treated for 1 hour. Subsequently, culture media was collected from donor group to replace the media of the receiving cells following a quick PBS wash. The receiving cells were then incubated for 45 minutes with the transferred media. Cells were then washed with PBS, fixed with 4% PFA, and blocked with blocking solution (2.5% normal goat serum and 2.5% normal donkey serum in PBST containing 0.2% Triton X-100). They were next incubated in an α-c-Myc antibody produced in rabbit (Sigma, C3956, 1:150) and an α-HA antibody produced in rat (MilliporeSigma, 11867423001, 1:150) at 4 °C overnight. After washing, cells were treated with appropriate secondary antibodies such as Alexa Fluor® 488 AffiniPure Donkey Anti-Rat IgG (H+L) (Jackson Immuno Research, 712-545-150, 1:1,000), Donkey α-rabbit IgG (H+L) Cy3^TM^ (Jackson Immuno Research, 711-165-152, 1:1,000), and Hoechst 33342 (Thermo Fisher Scientific, H3570, 1:20,000) at RT for 1 hour. Cells were then washed and mounted on the slide glass using fluromount-G® (SouthernBiotech, 011-01).

### Imaging

Immunocytochemistry and immunohistochemistry images were captured in a Zeiss 880 Airyscan confocal microscope with Zen Black software at the Fluorescence Microscopy Core Facility located at the University of Utah. For TEM, the JEOL JEM-1400 and the FEI Tecnai 12 transmission electron microscopes were employed. For SEM, FEI Quanta 600 FEG was used. The electron microscopes were located at the University of Utah’s Electron Microscopy Core Laboratory. All images were subjected to processing and analysis utilizing the open-source software Fiji, accessible at (https://fiji.sc/). Images concerning semithin sections were obtained using a Leica DM2500 optical microscope equipped with Leica Las software V3.8.

### CRISPR/Cas9 design and mutant mice generation

*TECTA^RS^* and *TECTA^ZP3^* mouse lines (C57Bl6/J background) were generated by CRISPR/Cas9-based genome editing at the University of Utah Transgenic and Gene Targeting Core. *TECTA* is located in an autosome (chromosome 9).

1. ***Tecta^RS^* mice:** For *Tecta^R2061S^* two sense and antisense guide RNAs (gRNA) targeting 5’-CTGAGCAGCAATAGCGCTGG-3’ (S2 targeting an upstream intron of E22) and 5’-CATGATGGACACACACCACA-3’ (S6 targeting the downstream intron of E22) were used. A 414 bp-long, reverse complementary single-strand oligo donor (ssODN) for *Tecta^RS^* was synthesized, comprising left-homology arm (75 bp), right-homology arm (75 bp), a mutation to change each protospacer adjacent motifs (PAM) in the intron, a point mutation to change Arg_2061_ (AGG) to Ser (AGC) and a silent mutation to introduce a unique EcoR1 (GAATTC) site. 2 μl of a mixture containing gRNA (30 ng/μl), the donor (50 ng/μl), and Cas9 protein (100 ng/μl) was injected per C57BL6/J pronuclear embryo.
2. ***Tecta^ZP3^* mice:** For *Tecta^ZP3^* a sense guide RNAs (gRNA) targeting 5’-ATGGAAGGAGCTGCAGAGGT-3’ (Tecta-E23-S6) was used to target the ω-site of the TECTA exon 23. A 260 bp-long, reverse complementary ssODN for *Tecta^ZP3^* was synthesized, comprising left-homology arm (56 bp), right-homology arm (58bp), the C-terminus of ZP3 (encoding aa 380-424: WTASAQTSVALGLGLATVAFLTLAAIVLAVTRKCHSSSYLVSLPQ) and an EcoR1 (GAATTC) site. 2 μl of a mixture containing gRNA (30 ng/μl), the donor (50 ng/μl), and Cas9 protein (100 ng/μl) was injected per C57BL6/J pronuclear embryo.

For both mice groups, 2 μl of a mixture comprising gRNAs, the donor, and Cas9 protein with a concentration at each 10 ng/μl, 2.5 ng/μl, and 25 ng/μl, respectively, was injected per C57BL6/J pronuclear embryo. After incubation overnight, injected embryos were implanted into the oviducts of pseudopregnant females. Genomic DNAs from G0 mice were used for PCR amplification. PCR products were cloned using TOPO®-TA cloning (Thermo Fisher Scientific, 450641), and were screened by EcoR1 digestion followed by Sanger sequencing. For TECTA^R2061S^ a 1,117 bp band was generated by RS-F (GCTAACAGCATTACGGGCCT) and RS-R (GCCGTTGTCTTCACACCAGT) primer sets. The overnight EcoR1 digestion for TECTA^R2061S^ yields 751 bp and 366 bp. Similarly, a primer set, 5’-CAGTGGGACCCATTAGGAG-3’ (DK76) and 5’-CTAGATGAAAGACCAGGGACC-3’ (DK77) generated a 748 bp band for TECTA^ZP3^, which was further cleaved by EcoR1 into 544 bp and 204 bp bands.

### Genotyping

A tail sample was collected from a P0 mouse pup and combined with 200 μl of 50 mM NaOH. It was then heated at 95 °C for 30 min, vortexed, mixed with 100 μl of 1 M Tris pH 6.8, and vortexed again. For subsequent analysis, 1 μl of the mixture was mixed with a PCR reagent kit (DreamTaq DNA Polymerase, Invitrogen, EP0703) according to the manufacturer’s instructions. The Tecta^R2061S^ generated a 1,117 bp band by RS-F (GCTAACAGCATTACGGGCCT) and RS-R (GCCGTTGTCTTCACACCAGT). The overnight EcoR1 digestion for TECTA^R2061S^ yields 751 bp and 366 bp. The TECTA^WT^ and TECTA^ZP3^ alleles generated a 606 bp band, and a 748 bp band by DK76 (CAGTGGGACCCATTAGGAG) and DK77 (CTAGATGAAAGACCAGGGACC) respectively. The PCR conditions are as follows: 94 °C for 30 sec, 60 °C for 30 sec, and 72 °C for 45 sec with 34 cycles. All the initial denaturing step and the final extension condition were 94 °C for 3 min and 72 °C for 10 min, respectively.

### Quantification and statistical analysis

All images were analyzed with the open-source software Fiji and the variations in all results were determined using GraphPad Prism9 software.

1. **Microvillus length:** We performed three replicates of each condition and five images per replicate were taken for quantification. The length of the microvilli was recorded along a horizontal line of 3 μm-length at the middle of the body domain. The microvillus length was averaged per image. The average length was used to perform the statistical testing. Since the variability of distribution was different between the groups (Brown-Forsythe test, p<0.0001), we performed Kruskal-Wallis non-parametric test with Dunn’s multiple comparison. Differences were considered statistically significant when p≤0.05.
2. **Microvillus density:** the number of the microvilli was recorded along a horizontal line of 3 μm-length at the middle of the body domain. Since the assumptions of ANOVA were fulfilled, we used an ordinary one-way ANOVA test to look for differences between the groups. A Tukey’s multiple comparison test was performed to assess the significance of differences between pairs of group means. Differences were considered statistically significant when p≤0.05.
3. **Collagen density:** We performed three replicates of each condition and five images per replicate were taken for quantification. The number of collagen fibrils was manually counted across a 1 μm line perpendicular to the collagen fibrils. Since the Kolmogorov-Smirnov test and the QQ plot of the measurements showed the normal distribution of the data, we performed an unpaired Student’s t-test. Differences were considered statistically significant when p≤0.05.
4. **Live cell surface labeling assay:** We performed three biological replicates of each condition, and five images per replicate were taken for quantification. The quantification of the co-localization of EPS8 and surface TECTA-ZP was performed by an independent researcher blind to the conditions. Each channel was thresholded to subtract the background signal, and the remaining pixels were selected as the region of interest (ROI). The ROI was then applied to the original images for measuring the integrated intensity (number of the pixels*average pixel intensity) of the surface TECTA-ZP signal in the ROI. The colocalization was measured as the percentage of the integrated intensity in the overlapped ROI over the total ROI and used for statistical analysis. Since the assumptions of ANOVA were fulfilled, we used an ordinary one-way ANOVA test to look for differences between the groups. A Tukey’s multiple comparison test was performed to assess the significance of differences between pairs of group means. Differences were considered statistically significant when p≤0.05.

## Results

### Ultrastructure of the developing TM and the apical surface of the TM-producing cells

The developing TM exhibits a higher-order ultrastructure composed of structurally distinct domains along the radial axis and multiple layers along the vertical axis (Figure 1A). To characterize the ultrastructure of collagen fibrils and non-collagenous fibers of the developing TM and their association with producing cells, we performed Airyscan high-resolution imaging of *Pisum sativum* agglutinin (PSA) lectin staining (Figure 1A), transmission electron microscopy (TEM) of the radial section of the mouse cochlea (Figure 1B-D), and scanning electron microscopy (SEM) (Figure 1E-F) of the cochlea at postnatal day (P) 2.

**Figure 1.**
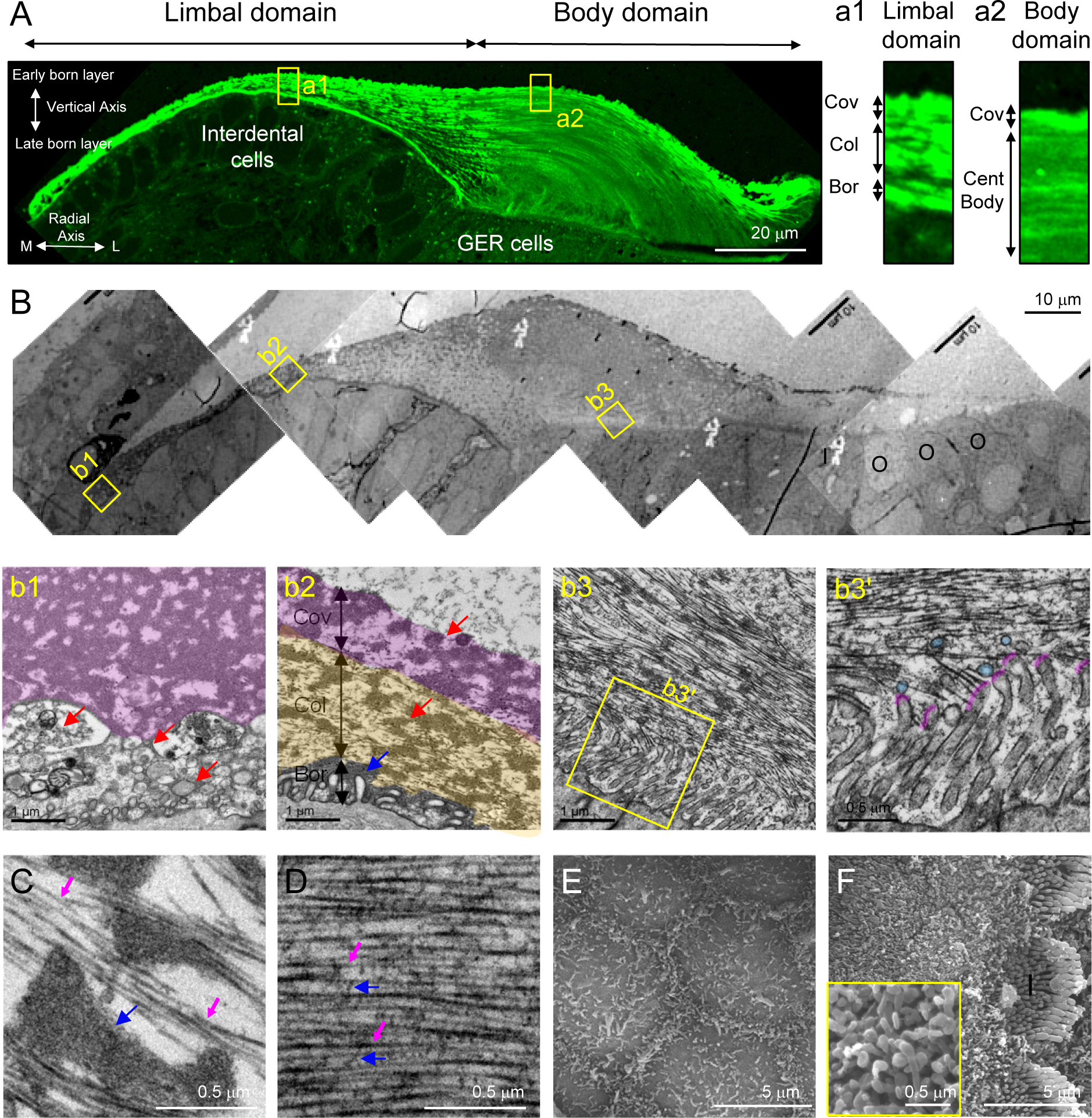
The shapes of the apical cell surface and the associated matrix are distinct between the limbal and body domains of the developing tectorial membrane (TM). A. PSA lectin staining of the TM at postnatal day (P) 2. The TM is composed of two domains along the radial axis: the limbal domain and the body domain. The limbal domain is produced by interdental cells and is composed of three layers: covernet (Cov), collagenous (Col), and border (Bor) layers. The developing body domain is produced by the greater epithelial ridge (GER) cells and is composed of two layers: covernet and central body (Cent Body) layers. PSA lectin staining shows distinct matrix architecture between the domains. M: medial; L: lateral. B. Transmission electron microscopy (TEM) of the TM at P2. **b1**: the most medial region of the limbal domain contains the extracellular vesicles (EVs; arrows) under the covernet layer (a pink area). **b2**: limbal domain. Dense non-collagenous fibers are present in all three layers of the limbal matrix (arrows). The apical surface of the interdental cells displays short and sparse microvilli. **b3** and **b3’**: body domain. The apical surface of the GER cells displays dense and long microvilli. Radially oriented collagen fibrils are associated with the microvillus tip of the GER cells (pink curves). EVs are present (marked in light blue) near the microvillus-ECM border. I: the inner hair cell; O: the outer hair cell. C. TEM of the limbal matrix. Dense non-collagenous fibers (blue arrows) are not associated with individual collagen fibrils. Collagen fibrils (pink arrows) are not associated with crosslinking fibers and are less ordered. D. TEM of the body matrix. Collagen fibrils (pink arrows) are associated with crosslinking fibers (blue arrows), evenly spaced, and parallelly organized. E. Scanning electron microscopy (SEM) of the interdental cells (P2). The TM is removed genetically from the spiral limbus (Otoa knockout mice). Hexagonally arranged interdental cells contain medial microvilli on the apical junction. The non-junctional apical membrane displays sparse microvilli. F. SEM of the GER cells. The TM is removed by force during the tissue prep. The apical surface of the GER columnar cells displays densely packed microvilli. I: the inner hair cell.

The interdental cells of the spiral limbus produced the limbal domain (Figure 1A). The limbal matrix was composed of three layers: covernet, collagenous, and border layers (Figure 1a1 and 1b2: Cov (purple), Col (yellow), and Bor (gray) layers, respectively). In this domain, the dense fibers composed of TECTA were observed in all three layers (Figure 1b2, arrows) (Andrade *et al*., 2016). The covernet layer facing the luminal side was the firstborn layer, covering the entire TM (Rueda *et al*., 2003) (Figure 1A). In the collagenous layer, dense tectorin fibers were clustered and separated from individual collagen fibrils (Figure 1b2, yellow areas, and 1C). The collagen fibrils in the limbal domain lacked crosslinking fibers (Figure 1C, magenta arrows) and were less ordered compared to those in the body domain (Figure 1D, magenta arrows). In the border layer, collagen fibrils were not directly associated with the apical surface membrane of the interdental cells. Instead, dense tectorin fibers were associated with the apical cell surface connecting the collagenous layer (Figure 1b2, a blue arrow). In the most medial region of the limbal domain, extracellular vesicles (EVs) were clustered between the covernet layer and the apical cell membrane (Figure 1b1, arrows). To visualize the apical surface of interdental cells, we used *Otoa* null mice, as their TM is detached from the spiral limbus (Zwaenepoel et al.,2002). SEM of the apical surface of the interdental cells showed a beehive-like junction with neighboring cells outlined by marginal microvilli (Cencer et al.,2023) (Figure 1E). The non-junctional apical membrane displayed a flat surface with sparse and short microvilli.

The greater epithelial ridge (GER) cells produce the body domain matrix (Figure 1A). The GER cells displayed dense microvilli on the apical surface (Figure 1b3 and 1F). The body matrix was composed of two layers during the growing phase: the covernet and the central body layer (Figure 1a2). Individual collagen fibrils in the body domain were interconnected with crosslinking fibers composed of TECTA and were evenly spaced (Figure 1b3’ and 1D, magenta and blue arrows mark collagen fibrils and collagen-crosslinking fibers, respectively) (Andrade *et al*., 2016). These parallel collagen fibrils were aligned radially along the medial-lateral axis (Figure 1b3). The border side end of collagen fibrils in this domain was attached to the microvillus tip of the GER cells (Figure 1b3’: marked in pink). EVs were observed within the matrix near the cellular border (Figure 1b3’ marked in light blue circles).

Overall, the ultrastructural analysis of the developing cochlea shows that the limbal and body domain-producing cells display distinct cell surface shapes, each associated with a unique matrix.

### TECTA processing

The surface expression of TECTA is required for collagen attachment to the microvillus tip membrane and is essential for the higher-order organization of the collagen fibrils (Kim *et al*., 2019). Thus, TECTA plays multiple roles in TM organization, forming the dense tectorin fibers in the covernet layer and the limbal domain, forming collagen-crosslinking fibers in the body domain, and mediating collagen attachment to the microvilli tip of the GER cells. TECTA contains the N-terminal NIDO domain, vWF domains, ZP domain, and GPI-anchorage signal (Figure 2A) (Legan *et al*., 2014). The NIDO and vWF domains mediate the interaction with collagenous and non-collagenous ECM proteins, respectively, and the ZP domain mediates the polymerization. The polymerization of the ZP domains is regulated by proteolytic shedding of the C-terminus on a consensus cleavage site (CCS). This cleavage removes the polymerization-blocking external hydrophobic patch (EHP) from the released ectodomain and induces the polymerization of the ZP domains in the extracellular space (Figure. S1A) (Jovine *et al*., 2005; Schaeffer et al.,2009). *In silico* analysis suggested that the ZP domain of TECTA (TECTA-ZP) may contain two potential CCSs (Lin et al.,2011; Bokhove et al.,2016) (Fig. S1A): (1) CCS1 contains a putative serine protease cleavage motif (XR↓IX) (Bokhove *et al*., 2016; Lucas et al.,2014) and (2) CCS2 contains a tetrabasic (RXXR↓) consensus Furin/proprotein convertase (PC) cleavage site (Lin *et al*., 2011).

**Figure 2.**
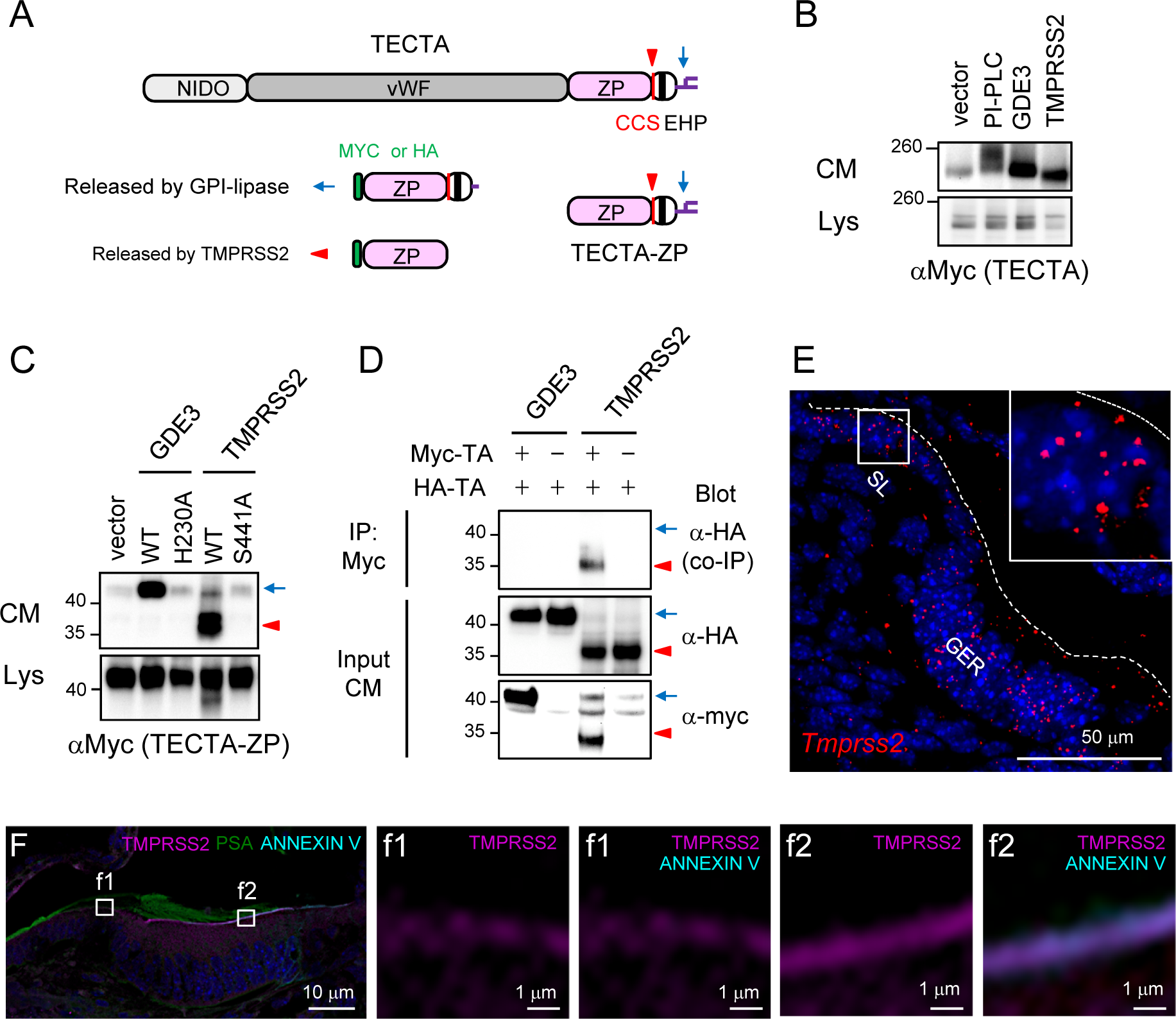
TMPRSS2 sheds TECTA. A. Diagram of TECTA and the ZP-domain of TECTA (TECTA-ZP). TECTA contains the NIDO domain, vWF domains, ZP domain, and C-terminal GPI-anchorage signal, which mediates collagen binding, ECM protein binding, polymerization, and GPI-anchorage, respectively. The ZP domain contains the consensus cleavage site (CCS) and the external hydrophobic patch (EHP) sequence close to the membrane-anchorage sequence. Cleavage of TMPRSS2 releases an ectodomain (red arrowheads), while cleavage by GPI-anchor lipases (blue arrows) releases non-proteolyzed protein. B. TECTA was expressed with an empty vector, GDE3 (a vertebrate GPI-anchor lipase) or TMPRSS2 in HEK293T cells, and the release of TECTA into the conditioned medium (CM) was monitored by western blot. The treatment of TECTA-expressing cells with phosphatidylinositol-phospholipase C (PI-PLC), a bacterial GPI-anchor lipase, or expression of GDE3 releases TECTA into the CM. TMPRSS2 releases a smaller fragment. Lys: lysate. C. Catalytic mutant of GDE3 (H230A) and TMPRSS2 (S441A) failed to release TECTA-ZP into the CM. *PNGaseF* was treated to both CM and Lys samples to show the size of the unglycosylated protein. D. Myc-TECTA-ZP and HA-TECTA-ZP were expressed with GDE3 or TMPRSS2. The released proteins in the CM were immunoprecipitated (IP) with Myc antibody and blotted with HA antibody. HA-TECTA-ZP released by TMPRSS2 was co-IPed by Myc antibody, while non-proteolyzed protein released by GDE3 was not. *PNGaseF* was treated to both CM and Lys samples to show the size of the unglycosylated protein. E. RNAscope in situ hybridization (ISH) of *Tmrpss2* in the developing cochlea at P0. Tmprss2 (red) is expressed in both the spiral limbus (SL) and GER. Hoechst (blue). F. Immunohistochemistry (ISH) of TMPRSS2 in the developing cochlear at P2. The TMPRSS2 puncta were observed in the cytoplasm and apical area of the interdental cells (f1) and GER cells (f2). TMPRSS2 antibody staining requires antigen retrieval, which reduces phalloidin signal (see materials and methods). Annexin V (cyan) is enriched on the microvilli (von der Mark and Mollenhauer,1997) and used to label the apical surface of the GER cells. PSA (green); Hoechst (blue).

To identify the cleavage site and the enzymes that mediate the release of TECTA, we performed a TECTA release assay from HEK293T cells. We overexpressed TECTA or TECTA-ZP in HEK293T cells with candidate enzymes and measured the release of TECTA into the culture medium by Western blot. As positive controls for TECTA release, we treated TECTA-expressing HEK293T cells with phosphatidylinositol-phospholipase C (PI-PLC), a bacterial GPI-anchor lipase, or co-expressed TECTA with GDE3, a vertebrate GPI-anchor lipase (Dobrowolski et al.,2020). As expected, TECTA was released into the medium by GPI-lipases, confirming the cell surface expression of GPI-anchored TECTA and TECTA-ZP (Figure 2B-D).

First, we tested the role of PC enzymes targeting CCS2 in TECTA processing. Overexpression of Furin or treatment of a pan-Furin/PC enzyme family inhibitor (decanoyl-RVKR-chloromethyl ketone: RVKR-cmK) (Tilak *et al*., 2014) did not affect the release of TECTA (not shown), indicating that the ZP domain of TECTA is unlikely a substrate of PC enzymes. Next, we tested whether TECTA is cleaved and released into the medium by the enzyme that can potentially target the CCS1. We tested transmembrane protease family members (TMPRSS-1, 2, 5, Prss36, Corin, and Ctsk) expressed in the developing cochlea (gEAR scRNAseq/Kelley(Kolla et al.,2020)) (not shown). Among the sheddases we tested, TMPRSS1 and 2 facilitated the release of TECTA into the culture medium (Figure S1B and 2B). TMPRSS2 robustly released TECTA, while Hepsin/HPN/TMPRSS1 released it at a lower level, suggesting that multiple enzymes may mediate the proteolytic shedding of TECTA at different degrees. The size of the released TECTA-ZP by TMPRSS2 was smaller than that of GPI-lipases and matched that of the truncated form at CCS1 (SR↓stop) (TECTA-ZP^Clv^, Figure S1B). Mutation of the catalytic amino acid (GDE3^H230A^ and TMPRSS2^S441A^) abolished the release, showing that the C-terminus of TECTA is the substrate of these enzymes (Figure 2C). Overall, our results showed that TMPRSS2 cleaves the ZP domain of TECTA at CCS1 and releases TECTA ectodomain *in vitro*.

### Proteolytic shedding of TECTA induces the multimerization of the ZP domains in the extracellular space

Next, we tested whether the proteolytic shedding impacts the multimerization of the released ectodomain in the extracellular space. We performed a homomeric co-immunoprecipitation (co-IP) assay between Myc- and HA-tagged proteins released by GDE3 (non-proteolyzed protein) or TMPRSS2 (proteolyzed ectodomain) from HEK293T cells (Fig. 2D). TECTA-ZP ectodomains shed by TMPRSS2 were co-IPed, while non-proteolyzed ZP domains released by GDE3 did not form a homomultimer. These results are consistent with previous observations that proteolytic shedding of the ZP domain facilitates the multimerization of the ZP domain-containing proteins in the extracellular space (Brunati et al.,2015). Since TECTA is organized into dense fibers in the covernet layer and limbal domain, we hypothesized that the proteolytic shedding TECTA may mediate the formation of dense tectorin fibers in these regions. This model further suggests that TECTA species released by different release modes ca\n be organized into distinct shapes.

We tested whether the expression of TECTA-cleaving enzymes is restricted to the cells that produce dense tectorin fibers. We examined the expression of *Tmprss1/Hpn* and *Tmprss2* by RNAscope *in situ* hybridization (ISH) at P0 (Figure 2E and S1C). Interestingly, these enzymes are expressed both in interdental cells and GER cells. Consistent with the ISH result, immunohistochemistry (IHC) showed that TMPRSS2 was expressed in both cell types (Figure 2F). TMPRSS2 puncta were detected in the cytoplasm and enriched on the apical area of these cells as marked by annexin V, which is present on the microvilli (von der Mark and Mollenhauer,1997). These observations indicate that the limbal domain-specific organization of dense tectorin fibers cannot be explained by the restricted expression of TECTA-shedding enzymes.

### Proteolytic shedding of TECTA is required for the formation of the dense tectorin fibers in the limbal domain and the covernet layer

While we were performing a mutagenesis screen targeting CCS1 to block TMPRSS2-dependent cleavage of TECTA, a deafness-associated mutation within CCS1 (*TECTA*: p.R2061S) was reported (Figure S1A) (Yasukawa et al.,2019). We hypothesized that TECTA^R2061S^ (TECTA^RS^) mutation in CCS1 may impact the proteolytic shedding of TECTA. Indeed, the release assay in HEK293T cells showed that the R2061S mutation dramatically reduced the shedding by TMPRSS2 (Figure 3A). The release by PI-PLC treatment or GDE3 expression was intact, indicating that the R2061S mutation did not affect the GPI-anchorage and surface expression of TECTA.

**Figure 3.**
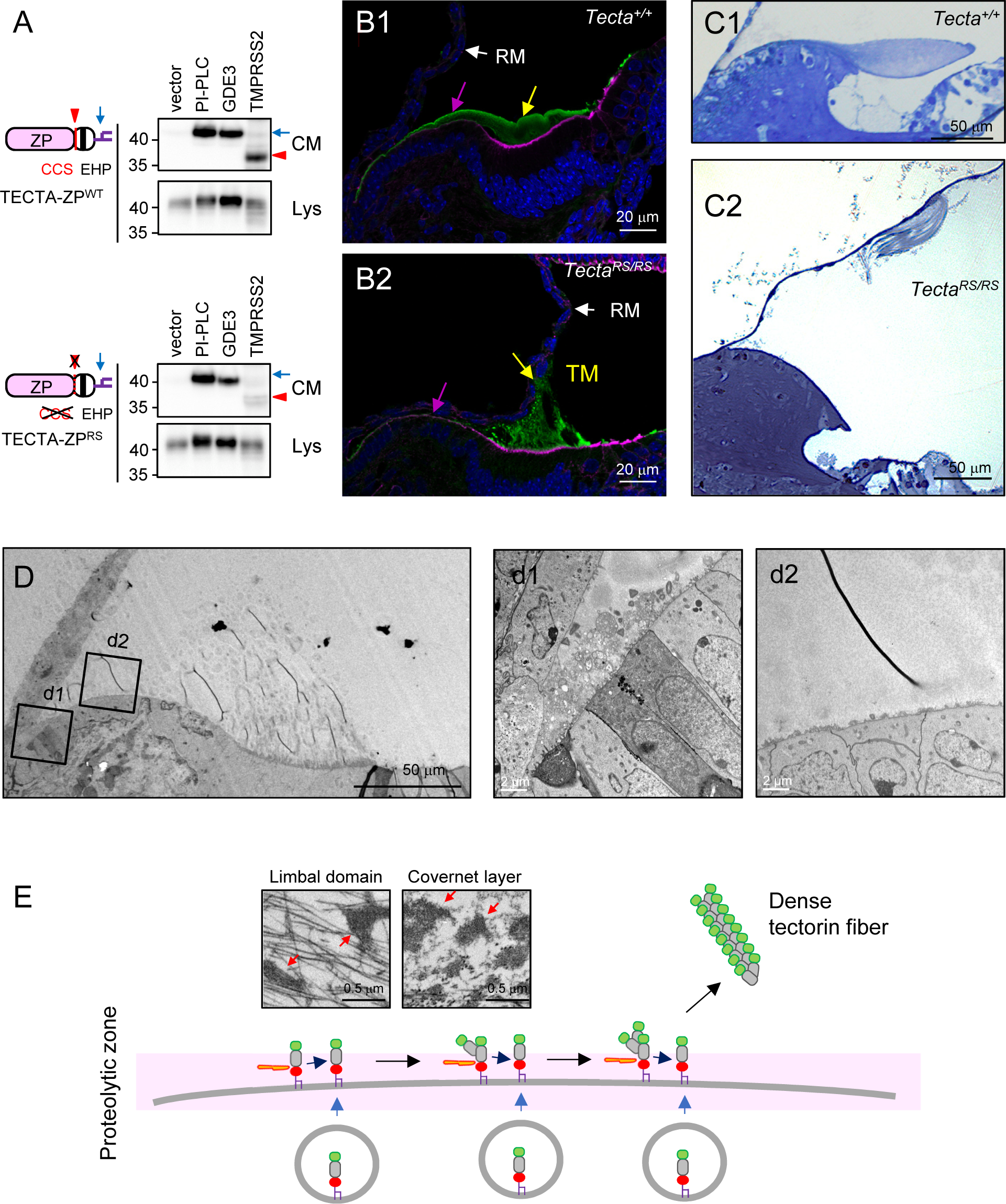
Proteolytic shedding of TECTA is required for the dense fiber formation in the limbal domain and the covernet layer. A. R2061S (RS) mutation on CCS1 reduced the release of TECTA-ZP by TMPRSS2, while the release by PI-PLC or GDE3 remained intact. B. Airyscan images of PSA (green) and phalloidin (magenta) at P2. The limbal domain (magenta arrows) and the covernet layer (yellow arrows) are absent in the TM of *Tecta^RS/RS^* mice. The body domain is associated with the GER cells but attached to the Reissner’s membrane (RM) in the absence of the covernet layer in *Tecta^RS/RS^* mice. Hoechst (blue). C. Semi-thin section followed by toluidine blue staining of the adult cochlea at 4 weeks. In wildtype mice (C1), the mature TM is associated with the spiral limbus of the cochlea. In *Tecta^RS/RS^* mice (C2), the TM is detached from the spiral limbus. D. TEM of the developing TM in *Tecta^RS/RS^* mice at P2. The dense fibers in the limbal domain is absent in *Tecta^RS/RS^* mice, resulting in the complete loss of the limbal domain (d2). The EV clusters in the most medium region remains intact (d1). E. Proposed model for formation of the dense tectorin fibers. Proteolytic shedding of TECTA is coupled with the elongation process of the ZP domain. Sequential process of the cleavage of the pre-assembled polymer and polymerization to the newly synthesized monomer will generate a long polymer species, which are stemming from the apical cell surface (3D printing model). The long polymer will form dense fiber in the limbal domain and covernet layer (red arrows).

Next, we generated a *Tecta^R2061S^* knock-in mouse line using CRISPR/Cas-9 to characterize the role of CCS1 cleavage of TECTA in TM organization *in vivo* (Figure S2). We first investigated the histology of the developing TM. PSA lectin staining of the TM at P2 showed that the covernet layer and limbal domain were absent in *Tecta^RS/RS^* mice (Figure 3B, yellow and magenta arrows, respectively). Notably, unlike other Tecta mutant lines where the TM is completely detached without surface-tethered TECTA (Legan *et al*., 2000; Kim *et al*., 2019), the body domain of *Tecta^RS/RS^* mice was assembled and attached to the GER cells. Interestingly, in the absence of the covernet layer, the body domain matrix was not radially oriented. Instead, it was vertically oriented and attached to the Reissner’s membrane (RM, Figure 3B, white arrows). This result revealed a role for the covernet layer: shielding the luminal end of collagen fibrils and preventing non-specific attachment.

Next, we characterized the structure of the mature TM by semi-thin sectioning followed by toluidine blue staining at 4 weeks (Figure 3C). In the wildtype, the mature TM remained associated with the spiral limbus, but the body domain was detached from the GER (Figure 3C1). However, in the absence of the limbal domain in *Tecta^RS/RS^* mice, the TM was detached from the spiral limbus and located in the luminal space (Figure 3C2). The body matrix was not completely disorganized but maintained a parallel architecture (Figure S2C).

TEM of the developing TM at P2 showed that the dense tectorin fibers were absent on the apical surface of the interdental cells in *Tecta^RS/RS^* mice, which resulted in the complete loss of the limbal domain (Figure 3D). Interestingly, the EVs clustered on the extreme medial region were preserved even in the absence of the limbal domain and the covernet layer (Figure 3d1).

Overall, these observations showed that proteolytic shedding of TECTA is necessary for the formation of dense tectorin fibers in the limbal matrix and the covernet layer, which are required for the attachment to the spiral limbus and radial orientation of the body domain matrix, respectively. These observations are consistent with the fiber formation process of ZP-containing proteins as implicated in a 3D printing model (Stsiapanava *et al*., 2020; Stanisich et al.,2020). In this model, the proteolytic shedding of a previously assembled polymer is coupled with the elongation process to a newly synthesized monomer. The repetition of this process results in a sequential polymerization of the fiber stemming from the cell surface (Figure 3E).

### Proteolytic shedding of TECTA determines the collagen-binding site in the body domain

If the proteolytic shedding of TECTA was occurring exclusively in the limbal domain to organize dense fibers, the body domain matrix should have remained intact in *Tecta^RS/RS^* mice. Surprisingly, TEM of the developing body matrix showed that the cell-ECM border was severely disorganized in the body domain of *Tecta^RS/RS^* mice at P2 (Figure 4). Notably, collagen fibrils were not attached to the tip membrane as observed in wildtype (Figure 4A, pink area). Instead, they were associated with the basal and lateral membrane of the microvilli (Figure 4B, an arrow). The average length of the microvilli was dramatically increased in *Tecta^RS/RS^* mice (Figure 4D). The length of the individual microvillus was variable, suggesting the instability of the microvillus architecture in the mutant mice (Figure 4E, p<0.0001, Brown-Forsythe test). The density of the microvilli remained intact (Figure 4F). These observations indicate that the specific attachment of collagen fibrils to the microvillus tip was not due to the steric hindrance of dense microvilli. Instead, the proteolytic cleavage of TECTA was required for the compartmentalization of the collagen-binding site. We hypothesized that the proteolytic shedding of TECTA may remove excessive surface TECTA on the basal and lateral membrane of the microvilli and restrict TECTA to the tip microdomain, which results in compartmentalization of the collagen-attaching microdomain.

**Figure 4.**
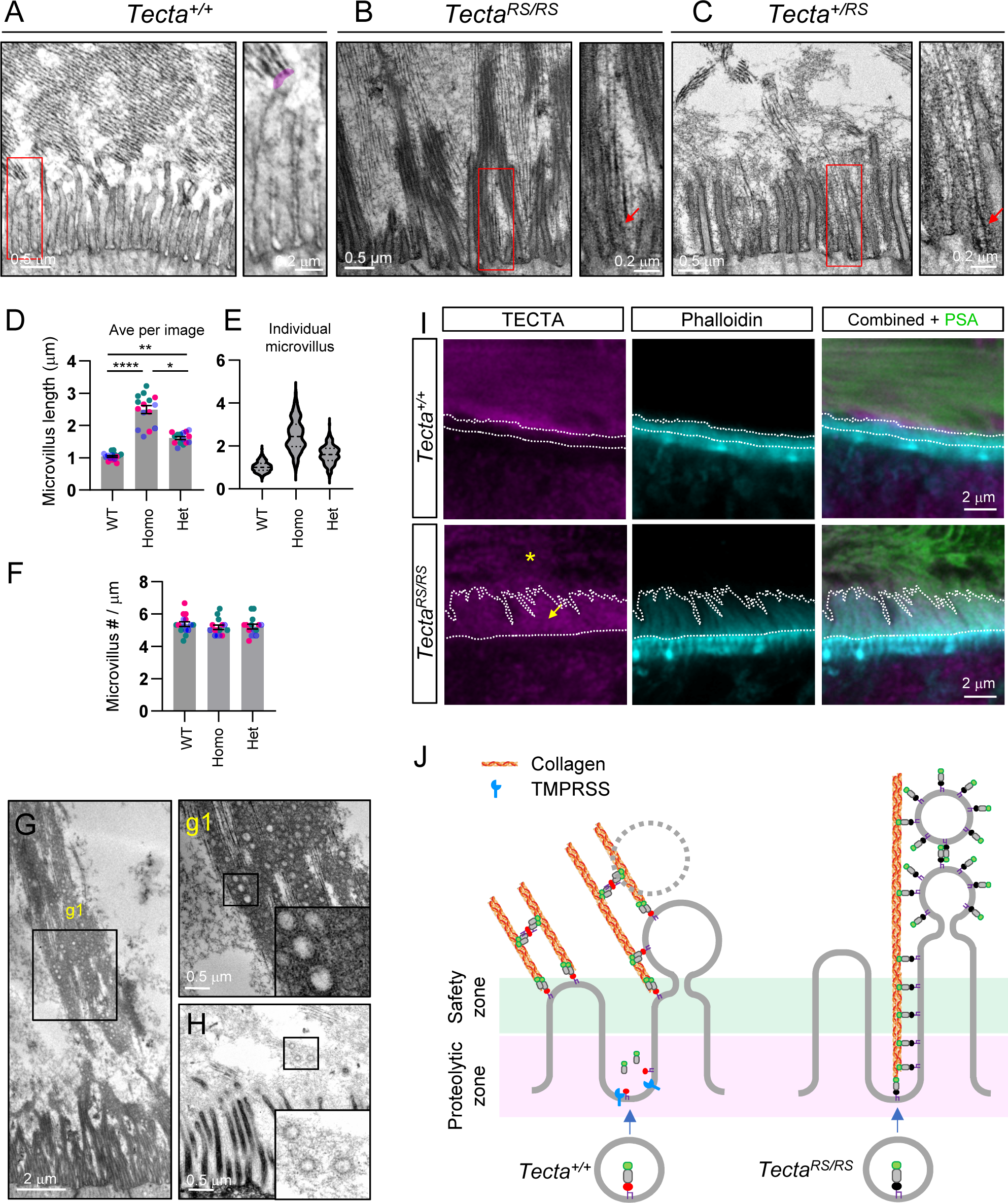
Proteolytic cleavage of TECTA compartmentalizes the collagen-attachment site on the microvilli in the body domain. A. TEM of the body domain of the wildtype at P2. Collagen fibrils are associated with the microvillus tip (marked in pink). B. TEM of *Tecta^RS/RS^* homozygous mice at P2. Collagen fibrils are associated with the basal and lateral membrane of the microvilli (an arrow). The length of the microvilli is increased and variable. C. TEM of *Tecta^+/RS^* heterozygous mice at P2. Collagen fibrils are associated with the basal and lateral membrane of the microvilli (an arrow). Intermediate fibers (dead-end polymer) stemming from the microvillus membrane. D. Quantification of the microvillus length. Left graph: an average of the microvillus length per image was plotted. N=3 animals per group, 5 images per animal. Data was plotted as mean with SEM. Each symbol represents an individual image from three animals (magenta, green, and blue). Adjusted ^∗∗∗∗^p < 0.0001; **p < 0.01: *p < 0.05 was significant by Kruskal-Wallis test, Dunn’s multiple comparisons. E. Right graph: the length of individual microvilli was plotted. N=3 animals per group, n=215-259 microvilli per group. Data was plotted in a violin plot outlining the kernel probability density with the width of the shaded area representing the proportion of the data located there. The median and quartiles are also represented on the graph. F. Quantification of the microvillus density. The number of the microvilli along the horizontal apical surface was plotted. N=3 animals, 5 images per animal. Data was plotted as mean with SEM. Each symbol represents an individual image from animals (magenta, green, and blue). Adjusted p value was not significant by one-way ANOVA, Tukey’s multiple comparisons. G. TEM of the body matrix above the cellular border *Tecta^RS/RS^* mice at P2. A large amount of EVs are accumulated in the lumen. H. TEM of the EVs in *Tecta^+/RS^* mice. EVs containing intermediated fibers are present near the microvilli. I. Airyscan IHC of TECTA showed that unlike wildtype TECTA, which is localized on the distal part of the microvilli (dashed curve), TECTA^RS^ is localized along the elongated microvilli. TECTA^RS^ is released from the cell and present within the matrix (an asterisk). J. Proposed model for the proteolytic cleavage of TECTA in the compartmentalization of the collagen-binding site. Proteolytic cleavage of TECTA removes excessive TECTA from the basal and lateral membrane of the microvillus. TECTA that reaches the microvillus tip may be protected from proteolytic shedding and recruit collagen fibrils. The EVs generated from the microvillus tip rapidly disintegrate. In *Tecta^RS/RS^* mice, the proteolysis-resistant form of TECTA is accumulated on the lateral membrane, which results in the ectopic association of the collagen fibrils. Dysregulation of the matrix-association domain may lead to the instability of the microvillus architecture and over-stability of the EVs in the luminal space.

We first investigated the TECTA protein localization by Airyscan high-resolution microscopy. In the wildtype TM, TECTA protein was accumulated on the distal part of the microvilli as marked by phalloidin staining. As previously observed, TECTA was released and localized within the matrix as marked by PSA staining (Fig 4I). However, in the body domain of *Tecta^RS/RS^* mice, TECTA^RS^ was strongly expressed on the lateral membrane of the elongated microvilli (Figure 4I, an arrow). TECTA^RS^ was also detected outside the cells (Figure 4I, an asterisk), indicating that there is an alternative release mechanism of TECTA. Overall, these observations show that the proteolytic shedding of TECTA regulates TECTA localization along the microvillus membrane.

To further test the hypothesis, we characterized the *Tecta^+/RS^* heterozygous mice to visualize the location of TECTA cleavage on the microvillus membrane. We reasoned that in the heterozygous mice, the incorporation of the cleavage-resistant form (TECTA^RS^) into the ZP polymer would block the elongation process, and the dead-end TECTA polymer would be tethered to the plasma membrane, marking the nearby TECTA cleavage site (Figure S3B1).

Indeed, we observed that intermediate fibers stemmed from the lateral membrane of the microvilli of *Tecta^+/RS^* mice (Figure 4C and S3B2), indicating that proteolytic cleavage of TECTA occurs on these sites. Collagen fibrils were mislocated to the basal and lateral membrane of the microvilli in *Tecta^+/RS^* mice, similar to *Tecta^RS/RS^* mice, further supporting that TECTA localization determines the collagen binding sites (Figure 4C, an arrow). The length of the microvilli in *Tecta^+/RS^* mice was between those of wildtype and *Tecta^RS/RS^* mice (Figure 4D-E). The covernet layer was formed in the heterozygote mice, albeit thinner than that of wildtype (Fig S3A-B).

These observations suggest that ectopic collagen attachment to the lateral side of the microvilli may lead to the instability of microvillus architecture and an increase in the microvillus length. In *Tecta^RS/RS^* mice, additional pulling force by vertical attachment of collagen fibrils to the Reissner’s membrane may contribute to a further increase in microvillus length. Interestingly, a large amount of EVs were accumulated in the luminal space of the *Tecta^RS/RS^* mice (Figure 4G). In *Tecta^+/RS^* mice, EVs covered with intermediate fibers (presumably a dead-end TECTA polymer) were observed (Figure 4H). Overall, our observations show that proteolytic shedding of TECTA in the body domain restricts TECTA to the microvillus tip, which results in the recruitment of collagen fibrils. Proper compartmentalization of the collagen-attachment site is critical for maintaining microvillus stability and EV homeostasis (Figure 4J).

### TECTA is compartmentalized to the microvillus tip by proteolytic cleavage

Next, we tested whether microvilli compartmentalize TECTA localization in a proteolysis-dependent manner. We used LS174T-W4 cells, a colon cancer cell line, of which microvilli were induced by doxycycline (DOX)-regulated activation of the polarity kinase LKB1 (Figure 5 and S4) (Hartmann *et al*., 2022) (Baas *et al*., 2004). We transfected TECTA-ZP with and without TMPRSS2 into the cells and performed cell surface staining of TECTA-ZP (sTECTA-ZP) two days after transfection. Confocal imaging showed that sTECTA-ZP puncta were evenly distributed on the plasma membrane without DOX treatment (Figure S4A). EPS8, a microvillus tip marker, was diffused in the cytoplasm in the absence of microvilli. Co-expression of TMPRSS2 did not impact the distribution pattern along the plasma membrane (Figure S4A).

**Figure 5.**
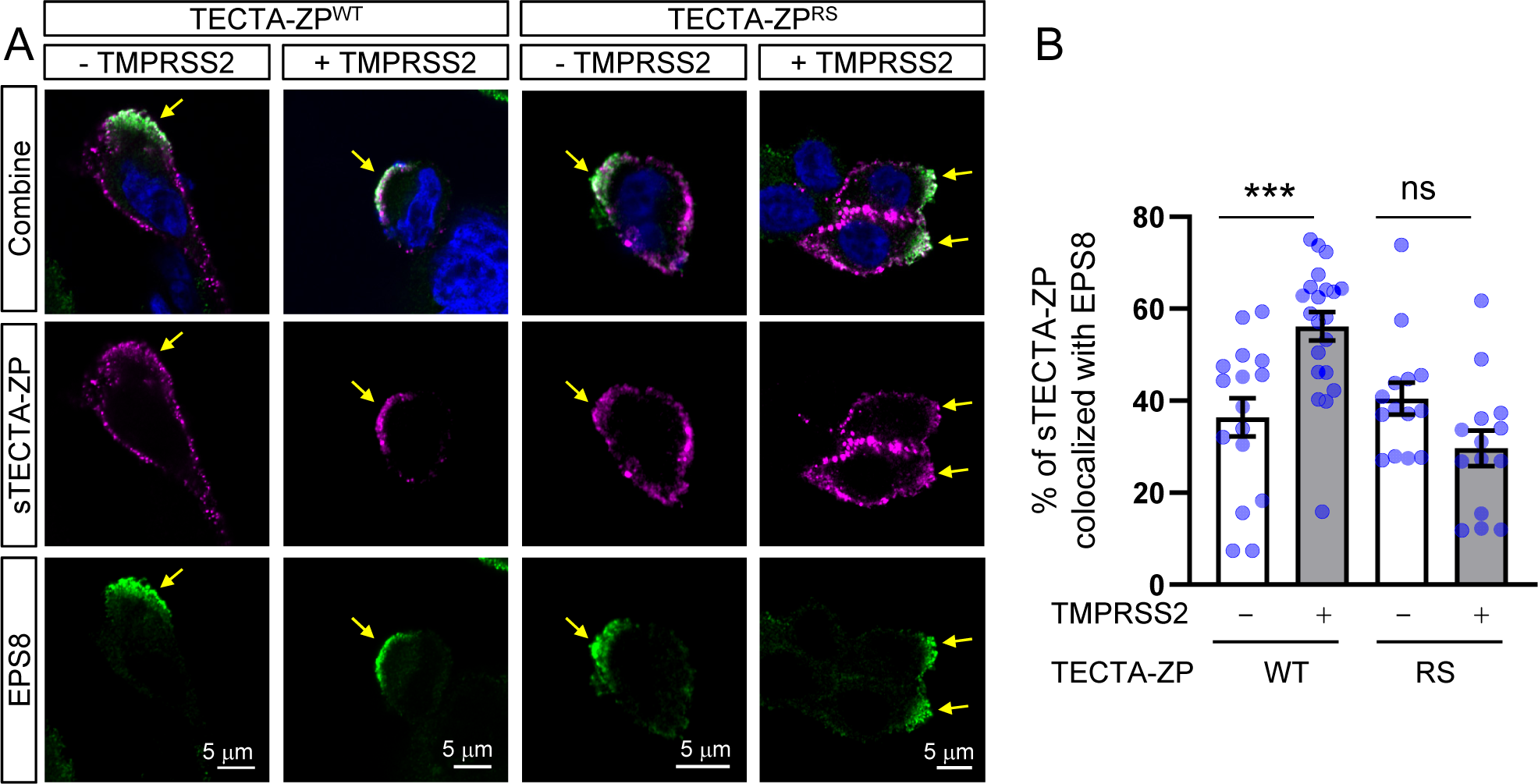
TECTA-ZP is enriched on the microvillus tip of W4 cells in the presence of TMPRSS2. A. LS174T-W4 cells (colon cancer cells) were transfected with TECTA-ZP with and without TMPRSS2 and the localization of the surface TECTA-ZP (sTECTA-ZP, magenta) and EPS8 (green), a microvillus tip marker, were measured. Doxycycline (DOX) treatment (1 μg/ml, 16 hours), which activates the polarity kinase LKB1, induces the accumulation of EPS8. sTECTA-ZP is co-localized with EPS8 in the presence of TMPRSS2 expression. TMPRSS2 expression did not induce the colocalization of the cleavage-resistant form (TECTA-ZP^RS^) with EPS8. Hoechst (blue). K. Quantification of the percentage of sTECTA-ZP colocalized with EPS8. The colocalization was measured as the percentage of the integrated intensity of sTECTA-ZP (number of the pixels*average pixel intensity) in the overlapped ROI over the total ROI. Data was plotted as mean with SEM. Each symbol represents an individual image from three pooled experiments (N=3, n=5 images per experimental condition). Adjusted ^∗∗∗^p < 0.001, or ns, not significant by one-way ANOVA, Tukey’s multiple comparisons.

DOX treatment for 16 hours induced the formation of the microvilli and the clustering of ESP8 on the microvillus tip (Figure S4B). Without TMPRSS2 overexpression, about 36±4.2% (mean±SEM, here and hereafter) of surface TECTA-ZP was co-localized with EPS8 puncta (Figure 5B). Notably, when TMPRSS2 was co-expressed, the percentage of sTECTA-ZP co-localized with EPS8 was significantly increased (Figure 5B, 56±3.1%, p=0.0008, one-way ANOVA, Tukey’s multiple comparisons). TMPRSS2 failed to induce the colocalization of EPS8 and TECTA-ZP^RS^, a cleavage-resistant form (40±3.5%, 30±3.9, without and with TMPRSS2, respectively. p=0.218). Overall, these observations show that both proteolytic shedding and microvillus architecture are required for the compartmentalization of surface TECTA.

If TECTA shed by TMPRSS2 outside the microvillus tip is diffused away from the cells, the level of remaining TECTA on the microvillus tip would be similar to that on the plasma membrane without TMPRSS2 overexpression. Interestingly, we observed an enrichment of sTECTA-ZP on the microvillus tip in the presence of TMPRSS2. Although it is difficult to compare the expression level between the cells after overexpression, this observation suggests that the TECTA ectodomain can be captured by surface-tethered TECTA. To test this, we performed a medium transfer assay (Figure S5). We transfected HEK293T cells (donor cells) with Myc-TECTA and TMPRSS2 and collected the conditioned medium. We applied the conditioned medium to receiving cells for 45 minutes and measured the accumulation of shed TECTA on the recipient cells by immunostaining with Myc antibody after fixation. TECTA expressed in the receiving cells (marked by HA) recruited transferred myc-TECTA. Recipient cells without overexpression of TECTA or cells treated with PIPLC to remove surface TECTA did not capture shed TECTA (Myc-tagged). These observations support the capturing of shed TECTA ectodomain by surface-tethered TECTA.

Overall, our *in vivo* and *in vitro* results show that the proteolytic shedding of TECTA removes excessive TECTA and restricts the localization/enrichment of TECTA on the microvillus tip, which specifies the collagen-association site (Figure 4J).

### GPI-anchorage of TECTA is required for the release of TECTA from the microvillus tip

How is TECTA released from the microvillus tip? Proteolysis-resistant TECTA^RS^ proteins were detected outside the cells, suggesting that there is an alternative release mechanism of TECTA (Figure 4I, an asterisk). Interestingly, a massive amount of EVs was present in the luminal space of *Tecta^RS/RS^* mice (Figure 4G), suggesting that TECTA^RS^ may be released from the microvilli via EVs. Microvilli are known to be an EV-producing organelle (McConnell and Tyska,2007; McConnell *et al*., 2009; Rilla,2021), and various sizes of EVs are produced from the microvillus tip of GER cells in the developing cochlea (Figure 1B3’ and 7A). In addition, GPI-anchored proteins are enriched on many forms of EVs, including microvesicles, exosomes, and nanoparticles (Sloand et al.,1998; Tutanov et al.,2023). Thus, we hypothesized that GPI-anchorage of TECTA may play a role in EV-dependent release of TECTA from the microvillus tip. To test this hypothesis, we generated a swapping mutant where normally GPI-anchored TECTA is converted into a transmembrane protein. We aimed to preserve the proteolytic shedding of the swapping mutant and chose the C-terminus of another ZP domain-containing protein, ZP3, of which the transmembrane domain is close to its CCS, and the cytoplasmic tail is short (Jimenez-Movilla and Dean,2011). Thus, the proximity of CCS1 to the membrane-tethering site, which can impact the substrate specificity of the transmembrane proteases, is likely preserved in the swapping mutant (Lichtenthaler et al.,2018) (Figure S6).

First, we tested whether TM domain swapping impacts the surface expression of TECTA *in vitro*. Surface biotinylation in HEK293T cells overexpressing TECTA^WT^ or TECTA^ZP3^ showed that a swapping mutation did not change the surface expression of the protein. As expected, PI-PLC treatment removed TECTA^WT^ from the cell surface but did not reduce the surface level of TECTA^ZP3^ (Figure 6B). TECTA-ZP^ZP3^ swapping did not affect the proteolytic shedding by TMPRSS2 (Figure 6C).

**Figure 6.**
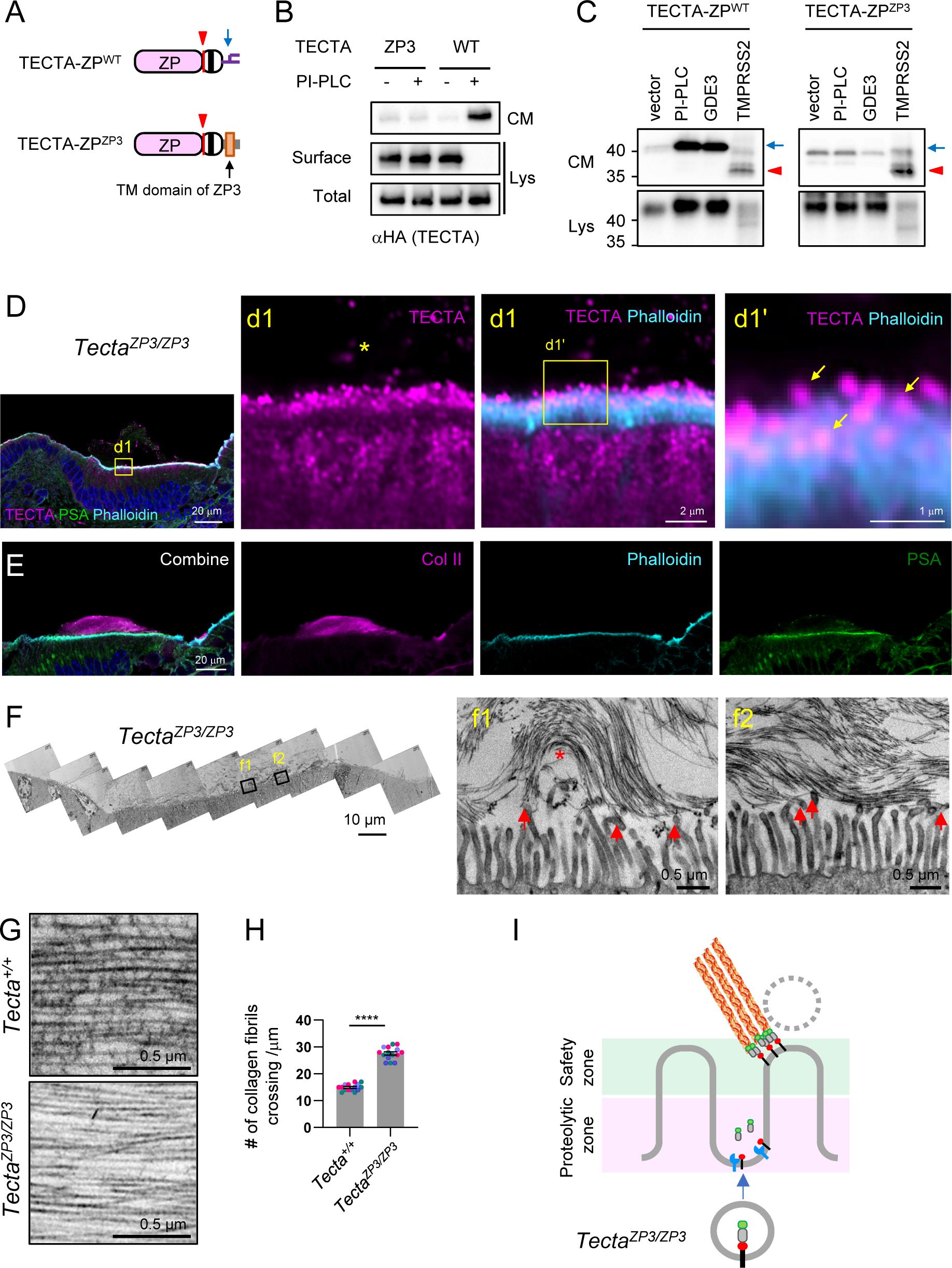
GPI-anchorage of TECTA is required for the release of TECTA from the microvillus tip. A. Diagram of the TECTA-ZP wildtype and swapping mutant with a transmembrane domain of ZP3. B. Surface biotinylation of TECTA-wildtype (WT) and TECTA-ZP3 swapping mutant expressed in HEK293T cells. TECTA-ZP was expressed on the cell surface membrane and not released by PI-PLC treatment, while TECTA-WT was removed from the cell surface and released into the CM. C. TMPRSS2 expression released TECTA-ZP^ZP3^ from HEK293T cells to the CM. GPI-lipases (PI-PLC treatment or GDE3 expression) did not induce the release of TECTA-_ZP_ZP3. D. Airyscan image of TECTA (magenta), phalloidin (cyan), PSA lectin (green), and Hoechst (blue) of the TM in *Tecta^ZP3/ZP3^* mice at P2. of TECTA-ZP^ZP3^ is localized on the microvillus tip (arrows). The release of TECTA-ZP^ZP3^ was dramatically reduced (an asterisk). E. Airyscan image of collagen II (Col2: magenta), phalloidin (cyan), PSA lectin (green), and Hoechst (blue) of the TM in *Tecta^ZP3/ZP3^* mice at P2. Collagen II is assembled in the TM and associated with the apical surface in *Tecta^ZP3/ZP3^* mice. F. TEM of the TM in *Tecta^ZP3/ZP3^* mice at P2. Collagen fibrils are associated with the microvillus tip of *Tecta^ZP3/ZP3^* mice (arrows). Collagen fibrils displaying a loop-like association at both ends were observed (an asterisk). G. TEM of the collagen fibrils of the body matrix. In wildtype mice, collagen fibrils are interconnected with collagen-crosslinking fibers and evenly spaced. Collagen fibrils of the *Tecta^ZP3/ZP3^* mice lack crosslinking fibers and are condensed. H. The number of crossing of collagen fibrils along the perpendicular line is increased in TectaZP3/ZP3 mice. Data are plotted as mean with SEM. The number of the collagen fibrils across a 1 μm line perpendicular to the fibers was manually counted and plotted. N=3 animals per group, 5 images per animal. Each symbol represents an individual image from three animals (magenta, green and blue). Adjusted ∗∗∗∗p < 0.0001, by unpaired Student’s t-test. I. Diagram of the TECTA localization and collagen arrangement in *Tecta^ZP3/ZP3^* mice. TECTA-ZP^ZP3^ is normally processed by proteases on the basal and lateral microvillus membrane and enriched on the microvillus tip. The tip-localized TECTA-ZP^ZP3^ recruits collagen fibrils. Without GPI-anchorage, TECTA is not released into the matrix, which results in the absence of collagen-crosslinking fibers. Collagen fibrils lacking crosslinking fibers are condensed and disorganized.

Next, we generated a *Tecta^ZP3^* allele by using CRISPR/Cas-9. Airyscan microscopy showed that unlike TECTA^RS^, which localized on the microvillus lateral membrane, TECTA^ZP3^ was highly enriched on the microvillus tip and barely detected on the lateral membrane of the microvilli (Figure 6d1’; arrows), indicating that the proteolytic shedding of TECTA that removes the excessive TECTA from the lateral membrane is intact in *Tecta^ZP3/ZP3^* mice. Interestingly, the release of TECTA^ZP3^ from the microvillus tip was dramatically reduced compared to that of wildtype and TECTA^RS^ (Figure 6d1: an asterisk), which resulted in reduced PSA lectin signal within the matrix. However, collagen fibrils were organized and associated with the apical membrane, as shown by Col2 staining (Figure 6E). These results indicate that the GPI-anchorage of TECTA is required for the release of TECTA from the microvillus tip.

Since TECTA^ZP3^ was highly accumulated on the microvillus tip, we expected the collagen fibrils to be associated with the microvillus tip in *Tecta^ZP3/ZP3^* mice. Indeed, TEM showed that collagen fibrils were exclusively associated with the microvillus tip in *Tecta^ZP3/ZP3^* mice (Figure 6F, arrows). Notably, collagen fibrils lacked crosslinking fibers, and the naked collagen fibrils were more densely packed than those associated with crosslinking fibers in the wildtype (Figure 6G-H). Collagen fibrils were less ordered, and a loop-shaped attachment where both ends of collagen fibrils were associated with the microvillus tip was also observed (Figure 6f1, an asterisk). These observations show that TECTA released from the microvillus tip organizes collagen-crosslinking fibers, which are critical for maintaining the spacing between collagen fibrils and a higher-order organization of the collagen matrix of the body domain.

### The potential role of EV in matrix organization

The EVs were produced from the microvillus tip of the GER cells (Figure 7A). The microvillus-derived EVs did not contain electron-dense materials within their lumen and appear to lack an actin cytoskeleton. Interestingly, the EVs were only observed on the border region of the matrix and not detected in the distal matrix away from the apical membrane in wildtype mice, suggesting that microvilli-derived EVs rapidly disintegrate upon their release in the developing TM. On the other hand, in *Tecta^RS/RS^* mice, a dramatic accumulation of EVs was observed in the luminal space (Figure 4G and 7B), suggesting that TECTA^RS^ on the EV membrane may stabilize the EV in the extracellular space.

**Figure 7.**
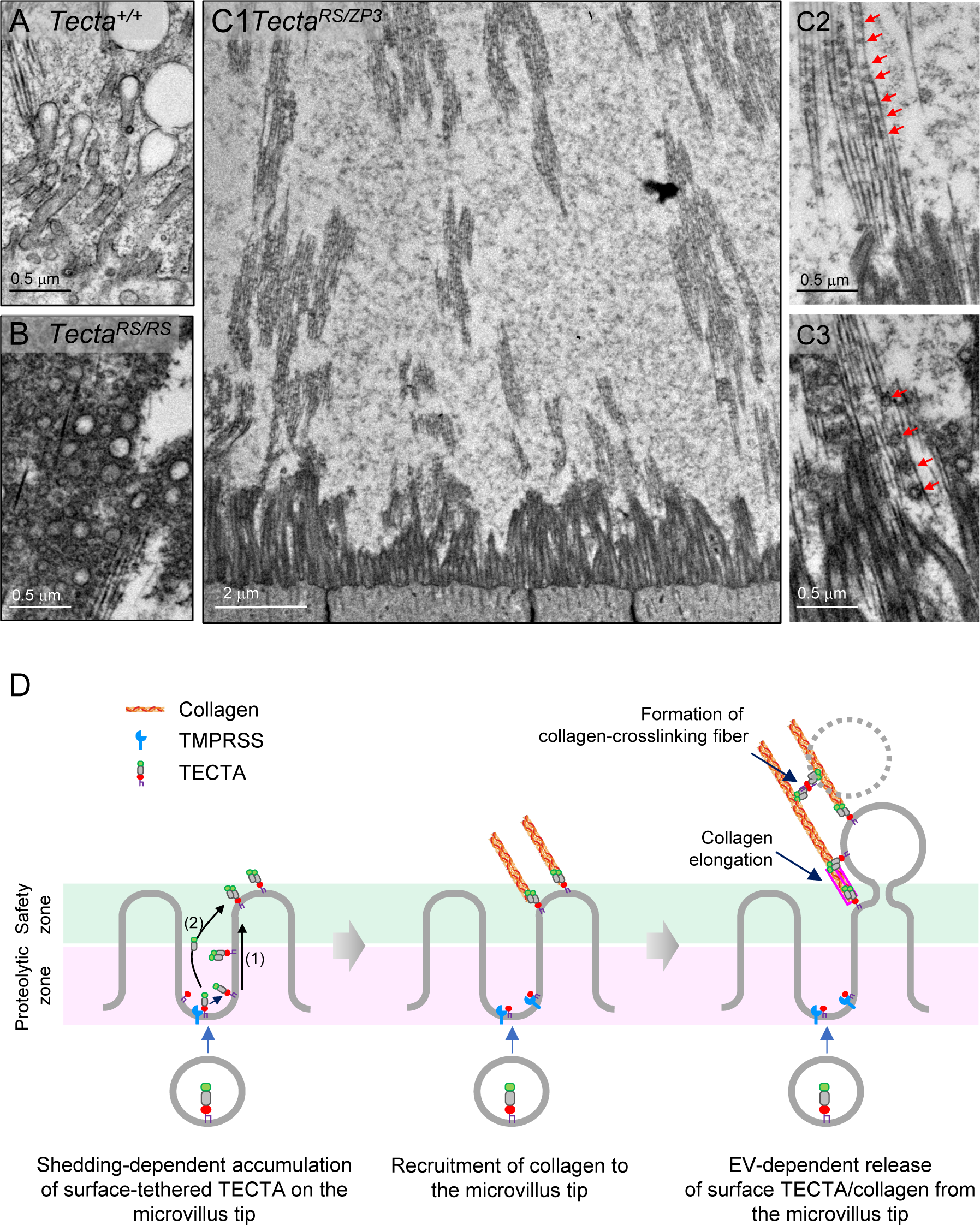
EVs are produced from the microvilli of the GER cells. A. TEM of the microvilli in wildtype mice at P2. Budding of the vesicles from the microvillus tip was observed. B. TEM of the lumen in *Tecta^RS/RS^* mice at P2. A large amount of EVs was accumulated in the lumen. C. TEM of the *Tecta^RS/ZP3^* mice at P2. EV-like circular clusters were observed along the collagen fibrils that are connected to the microvillus tip. D. Proposed model for the role of microvilli in collagen organization. Newly synthesized TECTA is delivered to the base of the microvilli and migrates to the microvillus tip along the membrane. Excessive collagen is removed from the basal and lateral membrane by proteolytic shedding. The shed TECTA ectodomain may form a dimer or a small oligomer with surface-tethered TECTA and migrate to the microvillus tip (1) or travel in the extracellular space and be captured on the microvillus tip (2). TECTA that reaches the microvillus tip (safe zone) is no longer processed and is not organized into a long polymer (dense tectorin fibers). The tip-localized TECTA recruits collagen fibrils. TECTA/collagen complex is released from the tip via EVs. The released EVs rapidly disintegrate, and dispersed TECTA forms the collagen-crosslinking fibers. The budding of EVs may expose the end of the collagen fibrils so a new collagen molecule (outlined in pink) is incorporated into the growing end.

Since TECTA^ZP3^ is strongly accumulated on the microvillus tip and maintains the attachment of collagen fibrils to the tip (Figure 6D-F), we reasoned we could visualize the EV loading events on the growing collagen fibrils in the presence of both TECTA^RS^ and TECTA^ZP3^ proteins. TECTA^ZP3^ can maintain the collagen’s attachment to the microvillus tip where EVs are produced, and TECTA^RS^ stabilizes EVs in the luminal space. Thus, we characterized the ultrastructure of the trans-heterozygous mice (*Tecta^RS/ZP3^* mice). Notably, the round-shaped electron-dense clusters, which appear to be the remnants of EVs, were observed along the collagen fibrils that were attached to the microvillus tip (Figure 3C). These observations suggest that the elongation of collagen fibrils and the formation of collagen-crosslinking fibers may be coupled with EV release. This coordinated process on the cell-matrix border may mediate the sequential outward growth of the higher-order apical ECM architecture from the cell surface (Figure 7D).

## Discussion

### Distinct release modes of TECTA

Our study shows that TECTA is released from the cell surface by two distinct mechanisms: proteolytic shedding and GPI-anchor-dependent release. The microvillus architecture spatially segregates two distinct release mechanisms, and each release mode is required to establish higher-order ECM architecture. In the limbal domain, where microvilli are sparse, TECTA is predominantly shed by proteolytic cleavage to form dense tectorin fibers (Figure 3E). In the body domain, where the apical cell surface displays dense microvilli, proteolytic shedding removes excessive TECTA from the basal and lateral membrane of the microvilli and restricts the surface-tethered TECTA to the microvillus tip membrane, which leads to the compartmentalization of the collagen-binding sites. TECTA on the microvillus tip is, in turn, released in a GPI-anchor-dependent manner to organize collagen-crosslinking fibers, which maintains the spacing and parallel organization of collagen fibrils (Figure 7D).

### Domain-specific organization

TECTA is expressed in both the limbal domain and body domain-producing cells, but forms distinct polymers. We originally hypothesized that proteolytic shedding, which is required to form dense fibers, might be restricted to the limbal domain. Proteolytic shedding occurs in the body domain as well, but plays a different role by restricting the surface-tethered TECTA to the microvillus tip (Figure 4C). These observations are consistent with the expression of Tmprss1 and 2 in both domains (Figure 2 and S1C). How is dense fiber formation by proteolytically shed TECTA inhibited in the body domain? (1) It can be mediated by distinct cell shapes. In the limbal domain, the apical surface of interdental cells contains sparse microvilli allowing the surface-tethered TECTA to be a substrate of the protease, whereas the dense microvillar protrusions of the GER columnar cells in the body domain may provide a safe zone that protects surface-tethered TECTA on the microvillus tip from proteolytic shedding (Figure 7D). The proteolytically shed TECTA from the lateral membrane can be delivered to the microvillus tip via two routes. (i) It can form a dimer or a small-size oligomer with GPI-anchored TECTA on the lateral membrane and migrate to the tip along the microvillus membrane. (ii) It can diffuse via the extracellular space and be captured by surface-tethered TECTA on the microvillus tip. Since the formation of dense fibers requires a continuous process of proteolytic shedding of multimers (Figure 3E), the TECTA oligomer on the microvillus tip will not be further processed into a long polymer (Figure 7D). The clustering of TECTA oligomers on the microvillus tip will generate multivalent binding sites for collagen, enabling a high-avidity association of collagen fibrils to the membrane. (2) It may be due to the different substrate selectivity of the protease activity between two domains. To generate a long polymer, the protease should preferentially cleave a polymer species instead of a newly synthesized monomer to maintain the elongation process of the polymer (Figure 3E). If the protease cleaves a monomer, the cleaved monomer would initiate a new polymerization process (dimer formation) or be diffused into the luminal space instead of being incorporated into the pre-assembled polymer. It is possible that the microenvironment of the membrane may differ between the two domains, which results in different substrate selectivity (polymer vs monomer) or catalytic efficiency, which may produce fibers with different lengths.

(3) Alternatively, it can be due to a domain-specific expression of other TM components. The TM is composed of collagenous (type II, V, IX, XI) and non-collagenous proteins, including otogelin (OTOG), OTOG-like, and carcinoembryonic antigen-related cell adhesion molecule 16 (CEACAM16), and beta-tectorin (TECTB) (Goodyear and Richardson,2018). While TECTA and Col II are expressed in both domains, some components display domain-specific expression patterns (https://umgear.org/) (Kolla *et al*., 2020). For example, the expression of Col IX is higher in the GER cells than in interdental cells. Col IX is a fibril-associated collagen with interrupted triple helices (FACIT) that generates protrusions on the collagen fibril that are associated with ECM molecules (Ricard-Blum,2011). The protrusion in the collagen fibrils of the body domain may recruit the proteolytically shed TECTA, preventing the formation of the dense fibers in the body matrix. The expression of TECTB and CEACAM16, which form striated sheets together with TECTA in adult TM, is higher in the GER cells. Formation of the striated sheet is impaired in the adult TM in *Tectb* knockout mice (Moreno-Pelayo *et al*., 2008). *Ceacam16* knockout mice show accelerated age-related deterioration of the TM (Goodyear et al.,2019). Although the gross architecture of the developing TM appears relatively normal in *Tectb* and *Ceacam16* knockout mice, the specific role of each component in the ultrastructure of each domain of the developing TM remains to be determined. Interestingly, in *Prdm16* cochlear-specific knockout mice, in which the maturation of the GER cells is delayed (fewer microvilli) and collagen expression is reduced, the body domain of the TM contains dense non-collagenous fibers as in the limbal domain, suggesting that the maturation status of the GER cells and balance between the expression level of collagens and TECTA may contribute to the matrix organization (a manuscript submitted). These possibilities are not mutually exclusive and may cooperate to build higher-order domain-specific architecture.

Interestingly, the dense fibers are absent in the limbal domain, and the covernet layer is thin in *Tecta^ZP3/ZP3^* mice (Figure 6D), while apical localization and TMPRSS2-dependent cleavage of TECTA^ZP3^ is intact (Figure 6C-D). These results indicate that the proteolytic shedding of TECTA is necessary but not sufficient for the organization of the dense fibers. The cytoplasmic tails of ZP3 prevent premature interaction with ZP2 within the cell (Jimenez-Movilla and Dean,2011). We cannot exclude the possibility that the cytoplasmic tail may differently regulate the protein-protein interaction of TECTA^ZP3^. GPI-anchorage of TECTA may be involved in the substrate selectivity of TMPRSS2 between polymer vs. monomer species. Further studies are needed to elucidate the role of GPI anchorage in the formation of dense tectorin fibers.

### TMPRSS and Hearing

TMPRSS1/Hepsin/Hpn mediates the shedding and polymerization of uromodulin, another ZP domain-containing protein expressed in kidney epithelial cells (Brunati *et al*., 2015). Interestingly, the TM is disorganized, and hearing is impaired in *Hpn* knockout mice (Guipponi et al.,2007). While the structure of the TM has not been investigated in *Tmprss2* knockout mice, the International Mouse Phenotyping Consortium (IMPC) reported that the threshold of the auditory brain stem response (ABR) is significantly elevated in *Tmprss2* knockout mice (https://www.mousephenotype.org/). These observations suggest that the TMPRSS family is involved in TECTA processing in the inner ear. While the TMPRSS family processes multiple substrates, it will be intriguing to investigate the specific role of each family member in the ultrastructural organization of the TM.

### Segregation of matrix-association and membrane-adhesion microdomains along the microvillus membrane

Many types of polarized epithelial cells display microvilli on their apical surface. They are often associated with apical ECM (Sharkova *et al*., 2022). Insect cuticles are associated with the tip of the microvilli of epidermal epithelial cells (Moussian,2013). Oocytes microvilli are surrounded by the zona pellucida matrix and mediate the communication between oocytes and somatic cells (Zhang et al.,2021a). In the gut epithelium, the microvilli are covered with a pericellular matrix, glycocalyx. Notably, ultrastructural analysis of the gut microvilli revealed that the matrix-association domain and membrane-adhesion domain are sharply segregated into the tip and lateral membrane of the microvillus, respectively (Sun *et al*., 2020). The matrix-association is mediated by the tip localization of mucin, and the membrane-association, called the intermicrovillar adhesion complex, is mediated by the cell adhesion molecules (Crawley *et al*., 2014). In the outer hair cells of the cochlea, the TM is connected to the tip of the tallest stereocilia (specialized microvilli-like protrusion), while membrane-adhesion complexes such as horizontal top connectors, shaft connectors, and ankle links are located to the lateral membrane of the stereocilia except for the tip links (Richardson and Petit,2019). Those observations indicate that compartmentalization of the microdomain along the microvillus membrane is tightly regulated. Our study shows that the segregation of the matrix-association domain and membrane-adhesion domain of the microvilli of the GER cells is mediated by proteolytic shedding of a matrix-binding protein (TECTA). Whether the compartmentalization of the matrix-association domain in other cells is mediated by the proteolytic shedding of matrix-interacting proteins remains to be determined. It would be intriguing to explore whether a similar segregation mode could be employed in the cells lacking microvilli. For instance, in neurons, the regulation of the microdomains for synaptic-adhesion proteins within the synaptic cleft and matrix-adhesion proteins in the perisynaptic regions may play a role in the regulation of synapse formation and stabilization (Dankovich and Rizzoli,2022).

### Maintenance of microvillus architecture by compartmentalizing the matrix-association domain

Intermicrovillar adhesion complex on the lateral membrane of a microvillus maintains the integrity of microvillus architecture. Genetic manipulations that disrupt the intermicrovillar complex result in short and irregular microvilli (Crawley *et al*., 2014; D’Alterio et al.,2005; Pinette et al.,2019). Interestingly, in *Tecta^RS/RS^* and *Tecta^+/RS^* mice, the microvilli are longer than in wildtype mice (Figure 4A-D). Collagen fibrils are associated with the lateral membrane of the microvilli of the GER cells in these mutant mice. This ectopic attachment of collagen fibrils may not only disrupt the intermicrovillar adhesion complex but also apply force to the attached microvilli. For example, elongation of the collagen fibrils, which are attached to the lateral membrane of the microvilli in these mice, may generate outward treadmilling force, which results in an increase in the microvilli length. In *Tecta^RS/RS^* mice, additional pulling force can be applied to the microvilli by the collagen fibrils that are vertically connected to Reissner’s membrane. These observations indicate that compartmentalizing the matrix-association and membrane-adhesion microdomains is critical for maintaining the integrity of the microvillus structure.

### Microvillus-derived EV in a matrix organization

Microvilli are known for EV-generating organelles (McConnell and Tyska,2007; McConnell *et al*., 2009). Microvillus-derived EVs are formed by the outward budding (called microvesicles) and are distinct from exosomes in their biosynthetic pathway, composition, and stability (Rilla,2021; Jeppesen et al.,2019; Nigro et al.,2021). Microvillus-derived EVs are known to deliver signaling molecules, including Hedgehog and cytokines, and regulate tissue development and immune response. (Hurbain *et al*., 2022; Kim *et al*., 2018).

We observed that microvesicles are produced from the tip of the microvillus during TM development. Interestingly, we observed that microvillus-derived EVs are only observed near the ECM-cellular border, not in the upper matrix, indicating that EVs derived from the microvillus tip rapidly disintegrate upon release. Unlike exosomes, microvesicles are selectively lost during sequential centrifugation steps, and their stability is highly sensitive to isolation procedures, indicating the unstable nature of microvesicles (Nigro *et al*., 2021). Notably, EVs are observed on the distal matrix in *Tecta^RS/RS^* or *Tecta^RS/ZP3^* mice, suggesting that the stability of microvesicles in the extracellular space can be determined by their membrane cargo. Zhang et al., recently reported that there is heterogeneity among the insoluble populations of EVs and nanoparticles (Zhang et al.,2021b). Interestingly, different GPI-anchored proteins are enriched in specific populations: GPI-anchored DPEP1 is enriched in exosomes, while GPI-anchored GPC1 is enriched in nanoparticles. While the biosynthetic mechanism of nanoparticles is not fully understood, it will be intriguing to test whether a population of nanoparticles enriched with apically localized GPI-anchored proteins is derived from the disintegration of microvillus-derived EVs or other microvesicles.

In *Tecta^RS/RS^* mice, a substantial quantity of EVs is accumulated in the extracellular space. It can be due to increased production and/or decreased disintegration of the EVs. The mislocalization of the TECTA^RS^ and ectopic collagen attachment on the lateral membrane may cause membrane instability of the microvilli, which leads to the overproduction of EVs (Shurer et al.,2019). However, in *Tecta^+/RS^* mice, EVs do not accumulate in the extracellular space despite similar mislocalization of TECTA and collagen attachment. Distinct multimeric states of TECTA [monomer in *Tecta^RS/RS^* mice (Figure 7C) and dead-end polymer in *Tecta^+/RS^* mice (Figure 4SB)] may cause different degrees of instability of the microvillus membrane and stability of the EV membrane.

### Extracellular assembly vs 3D printing model for the organization of the ECM

We previously showed that cell surface expression and release of TECTA are required for TM organization and proposed a 3D printing model for the organization of the apical ECM. There are two models for ECM organization: the extracellular assembly model and the 3D printing model. Many ECM components, including collagens and laminins, have a self-assembly activity. The ECMs that are organized within a confined space, such as the basement membrane and interstitial matrix, would be assembled predominantly in the extracellular assembly mode. Subsequent interaction of the assembled ECM with the cell surface receptors will further shape the matrix architecture. On the other hand, an apical ECM, which is organized in a large luminal space or facing the outside of the body, requires membrane-tethering of the growing matrix and is generated in an inside-out fashion (old matrix outside and newly synthesized matrix inside). In the present study, we propose that the 3D printing model can be applied to both the organization of the dense tectorin fibers in the limbal domain and larger-scale collagen elongation in the body domain. In the limbal domain, coupling of the proteolytic shedding of TECTA and ZP domain polymerization will lead to the sequential elongation of a tectorin fiber stemming from the plasma membrane (Figure 3E), as proposed in the uromodulin polymerization (Stsiapanava *et al*., 2020; Stanisich *et al*., 2020). In the body domain, the inside-out growth of the matrix can occur on a larger scale: elongation of collagen fibrils and incorporation of collagen-crosslinking fibers from the border side. Evenly spaced EV-like clusters are observed along the collagen fibrils when EVs are stabilized in *Tecta^RS/ZP3^* mice, suggesting that EV release events are orchestrated with collagen elongation. We propose that microvillus-derived EVs play dual roles in the organization of the TM. First, EVs release the TECTA-collagen complex from the microvillus tip. The released TECTA forms collagen-crosslinking fibers after the disintegration of EVs. Second, the budding of the membrane from the tip would expose the end of the collagen fibril and provide a space so a new collagen molecule can be incorporated into the growing collagen fibril (Figure 7D). This process may enable collagen fibrils to elongate while remaining associated with the apical membrane. This model predicts the elongation of the growing collagen molecules from the border side. Further studies are needed to determine where a newly synthesized collagen molecule is incorporated during the growing phase of the apical ECM *in vivo*.

Overall, our study showed that different release modes of a matrix-binding protein result in a distinct matrix shape of the apical ECM, and orchestrating these release modes is required to build higher-order ECM architecture. The shape of the apical cell surface plays a critical role in the morphogenesis of the associated ECM.

## Acknowledgement

We are grateful to Drs. Jason Shepherd (University of Utah), Monica Vetter (University of Utah), and Richard Cheney (University of North Carolina at Chapel Hill) for discussion and assistance. We are grateful to Drs. Willem-Jan Pannekoek (Center for Molecular Medicine, Utrecht, The Netherlands) for LS174T-W4 cells. We are grateful to Dr. Jan Christian (University of Utah) for providing Furin and BMP4 constructs and a pan-PC inhibitor. This work was supported by R21DC016750 and R01DC018814 to S.P.

