## Supplemental file for "Microvilli regulate the release modes of alpha-tectorin to organize the domain-specific matrix architecture of the tectorial membrane"

Figure S1

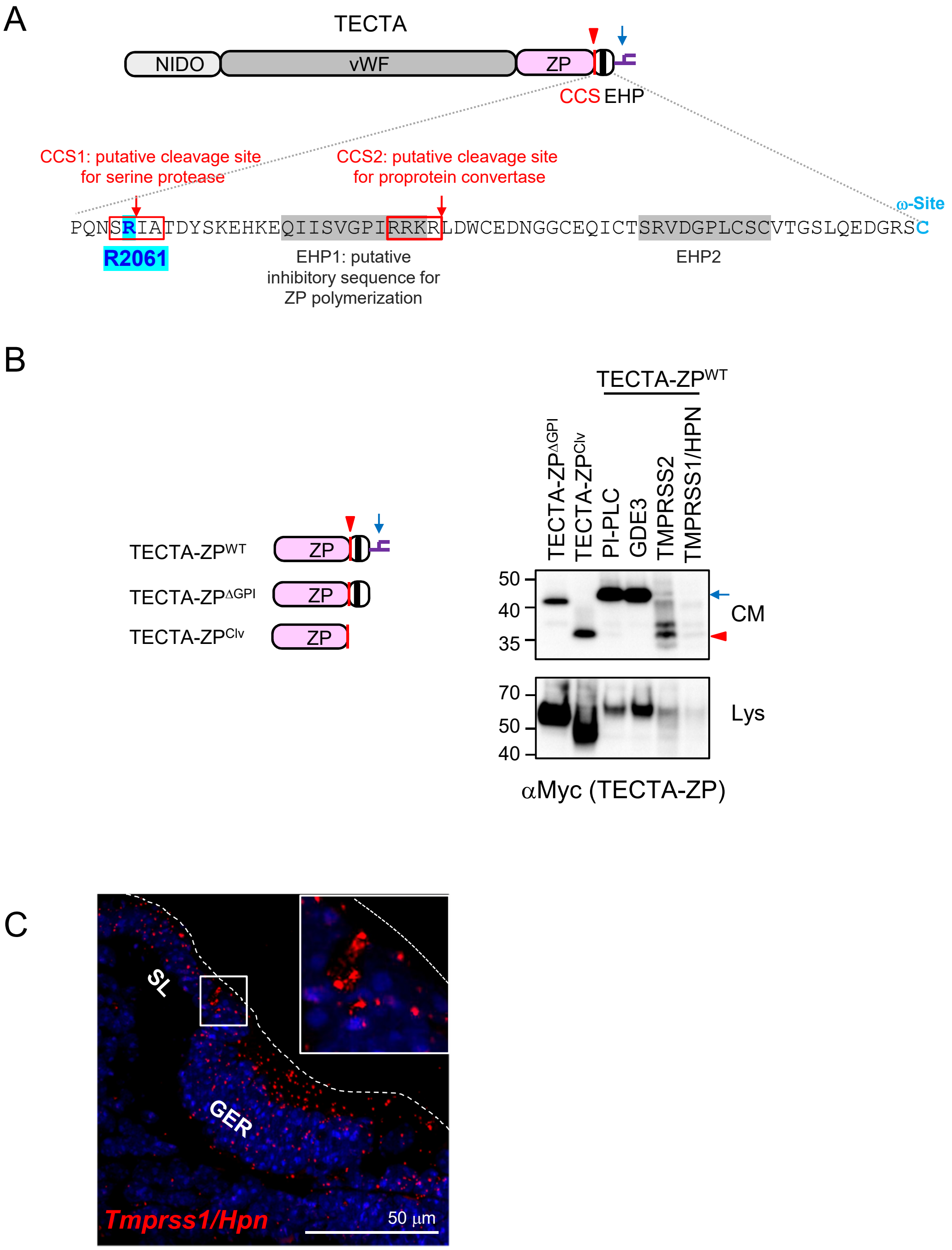
A
TECTA
NIDO
vWF
ZP
CCS
EHP
CCS1: putative cleavage site for serine protease
CCS2: putative cleavage site for proprotein convertase
ω-Site
PQNSRIATDYSKEHKEQIISVGPIRRKRLDWCEDNGGCEQICTSRVDGPLCSCVTGSLQEDGRSC
R2061
EHP1: putative inhibitory sequence for ZP polymerization
EHP2
B
TECTA-ZPWT
TECTA-ZPΔGPI
TECTA-ZPClv
ZP
TECTA-ZPΔGPI
TECTA-ZPClv
PI-PLC
GDE3
TMPRSS2
TMPRSS1/HPN
50
40
35
CM
70
50
40
Lys
αMyc (TECTA-ZP)
C
SL
GER
Tmprss1/Hpn
50 µm

**Figure S1. Related to Figure 2.**

A. The C-terminal sequence of the ZP domain of TECTA. *In silico* analysis suggested that there may be two putative CCS and EHP pairs: CCS1 for the serine protease cleavage site and CCS2 for the Furin/proprotein convertase cleavage site. Red arrows mark the cleavage site. The location of R2061 (mutated in CCS1) and omega ($\omega$)-site is marked in blue.

B. TMPRSS2 and TMPRSS1/HPN released the same size band of TECTA-ZP wildtype (TECTA-ZP$^{Clv}$) as the truncated form on CCS1 (TECTA-ZP$^{Clv}$: SR-stop) (a red arrowhead), while GPI-anchor lipases released the similar size band as the secreted form without the GPI-anchorage signal (TECTA-ZP$^{\Delta GPI}$). *PNGaseF* was treated to the CM to show the size of the unglycosylated protein.

C. RNAscope *in situ* hybridization (ISH) of *Tmrpss1/Hpn* in the developing cochlea at P0. *Tmprss1* (red) is expressed in both the spiral limbus and GER. Hoechst (blue).

Figure S2

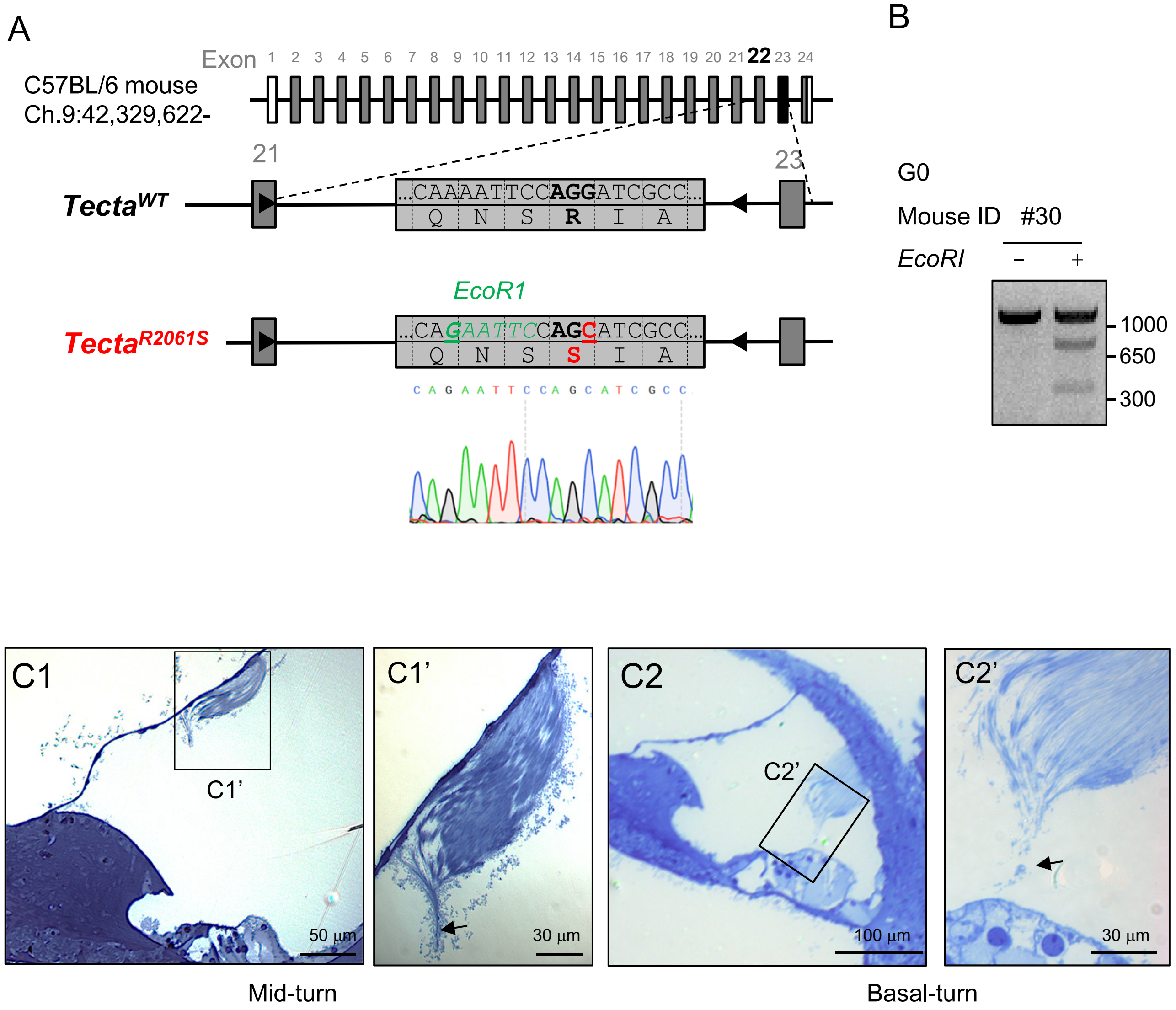
A
B
Exon 1 2 3 4 5 6 7 8 9 10 11 12 13 14 15 16 17 18 19 20 21 22 23 24
C57BL/6 mouse
Ch.9:42,329,622-
21
23
TectaWT
...CAAAATTCCAGGATCGCC...
Q N S R I A
EcoR1
TectaR2061S
...CAGAATTCCAGCATCGCC...
Q N S S I A
C A G A A T T C C A G C A T C G C C
G0
Mouse ID #30
EcoRI
– +
1000
650
300
C1
C1'
50 µm
30 µm
C2
C2'
100 µm
Mid-turn
Basal-turn

**Figure S2. Related to Figure 3.**

A. Generation of *Tecta*$^{RS/RS}$ mice by CRISPR/Cas-9. p.R2061 was mutated to S on exon 22. A silent mutation on p.Q2058 was introduced to generate an *EcoRI* site. Sanger sequencing confirmed the targeted mutation.
B. PCR followed by EcoRI treatment showed the targeted mutation in the G0 founders.
C. Semi-thin section followed by toluidine blue staining of adult cochlea at 4 weeks (related to Figure3C). Enlargement of the area in Figure 3C (C1') and the basal turn of the TM (C2') showed that the parallel organization of the matrix is maintained even in the detached TM. The attachment to the hair cell stereocilia appeared to be maintained as marked by a trace pointed to the outer hair cells (arrows).

Figure S3

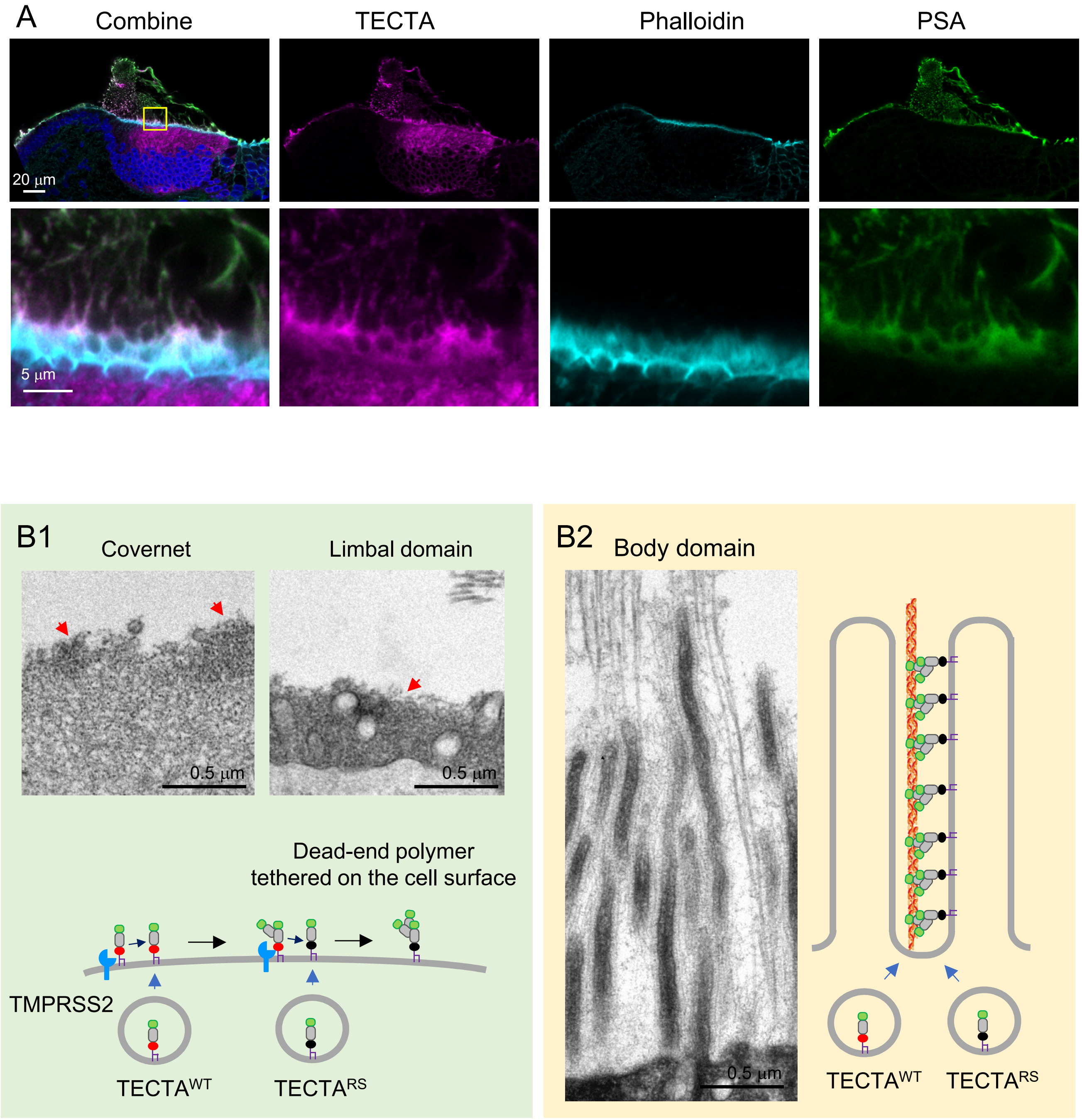
A
Combine
TECTA
Phalloidin
PSA
20 µm
5 µm
B1
Covernet
Limbal domain
0.5 µm
0.5 µm
Dead-end polymer
tethered on the cell surface
TMPRSS2
TECTA^WT
TECTA^RS
B2
Body domain
0.5 µm
TECTA^WT
TECTA^RS

**Figure S3. Related to Figure 4.**

A. Airyscan image of TECTA (magenta), phalloidin (cyan), PSA lectin (green), and Hoechst (blue) of the TM in *Tecta*$^{+/RS}$ heterozygous mice at P2. The thin limbal domain and covernet layer are formed in the heterozygous mice. TECTA is present along the microvilli and within the matrix.
B. TEM of the covernet layer and limbal domain (B1) and body domain (B2) of *Tecta*$^{+/RS}$ heterozygous mice. Incorporation of the mutant protein into the tectorin fiber will generate a dead-end polymer, which is tethered on the cell surface.

Figure S4

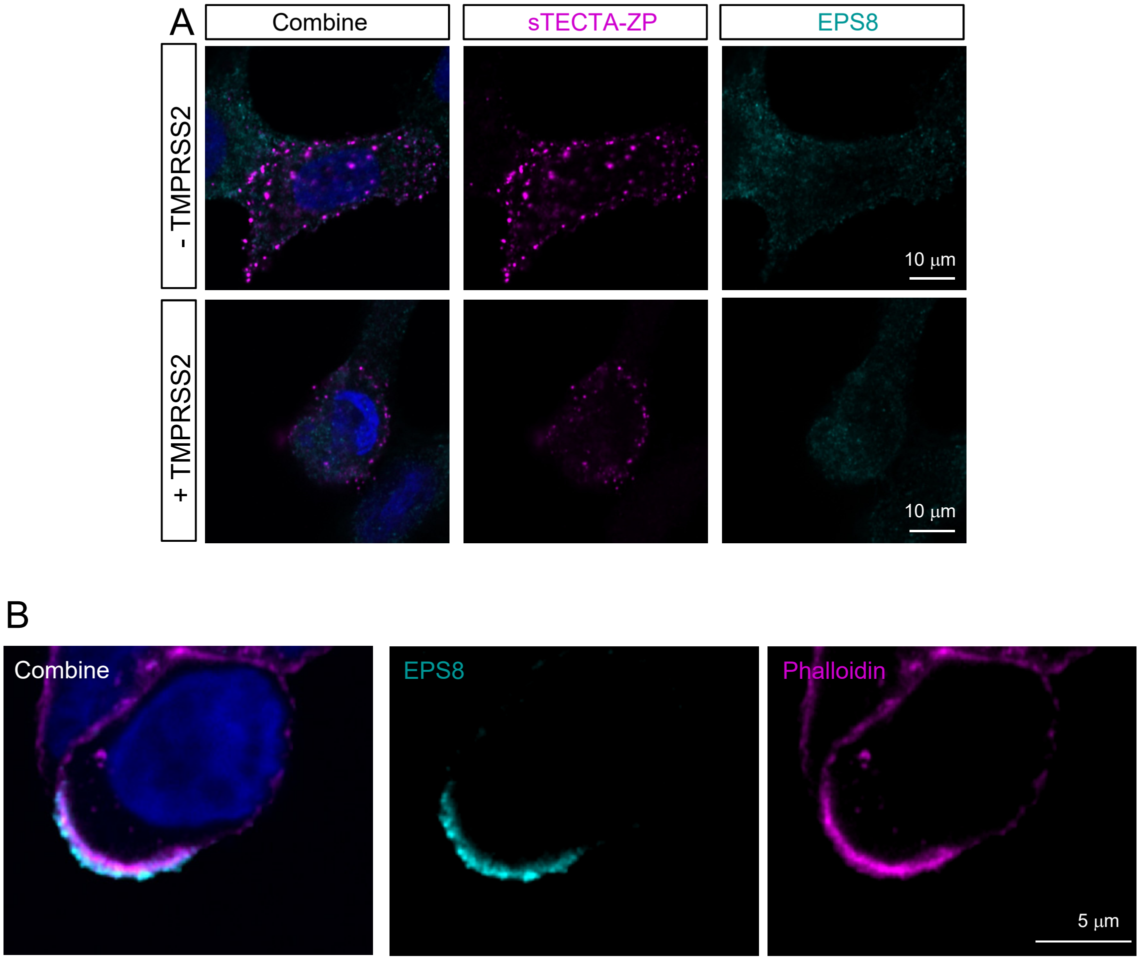
A
Combine
sTECTA-ZP
EPS8
- TMPRSS2
+ TMPRSS2
10 μm
10 μm
B
Combine
EPS8
Phalloidin
5 μm

**Figure S4. Related to Figure 5**

A. EPS8 is not accumulated in the absence of DOX treatment and diffused in the cytoplasm. sTECTA-ZP (magenta) is punctated on the cell surface. TMPRSS2 expression did not affect the distribution pattern of sTECTA-ZP without DOX treatment. EPS8 (cyan); Hoechst (blue).

B. EPS8 (cyan) is accumulated on the microvillus tip after DOX treatment (1 μg/ml, 16 hours). Phalloidin (magenta); Hoechst (blue).

Figure S5

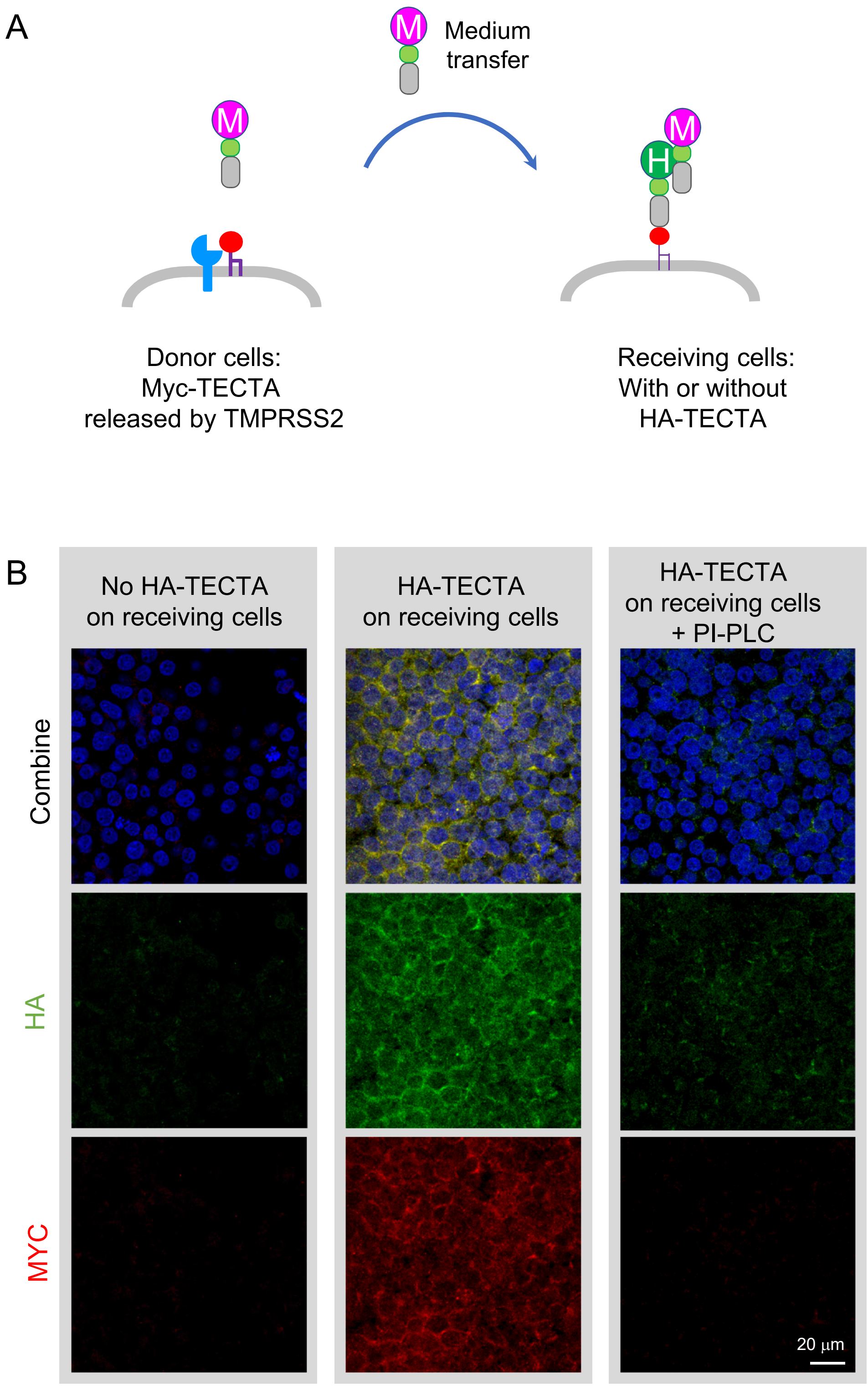
A
M
Medium
transfer
M
M
H
Donor cells:
Myc-TECTA
released by TMPRSS2
Receiving cells:
With or without
HA-TECTA
B
No HA-TECTA
on receiving cells
HA-TECTA
on receiving cells
HA-TECTA
on receiving cells
+ PI-PLC
Combine
HA
MYC
20 µm

**Figure S5. HEK293T cells transfer assay**

A. The conditioned medium of HEK293T cells transfected with Myc-TECTA and TMPRSS2 (donor cells) is applied to the receiving cells for 45 minutes. After the washout of the conditioned medium, the recruitment of Myc-TECTA on the receiving cells was determined by Myc antibody staining after fixation.

B. HEK293T cells without TECTA did not recruit Myc-TECTA (red). Expression of HA-TECTA (green) in the receiving cells facilitates the recruitment of Myc-TECTA. Removal of the surface HA-TECTA from the receiving cells by PI-PLC treatment reduced the recruitment of the Myc-TECTA. Hoechst (blue)

Figure S6

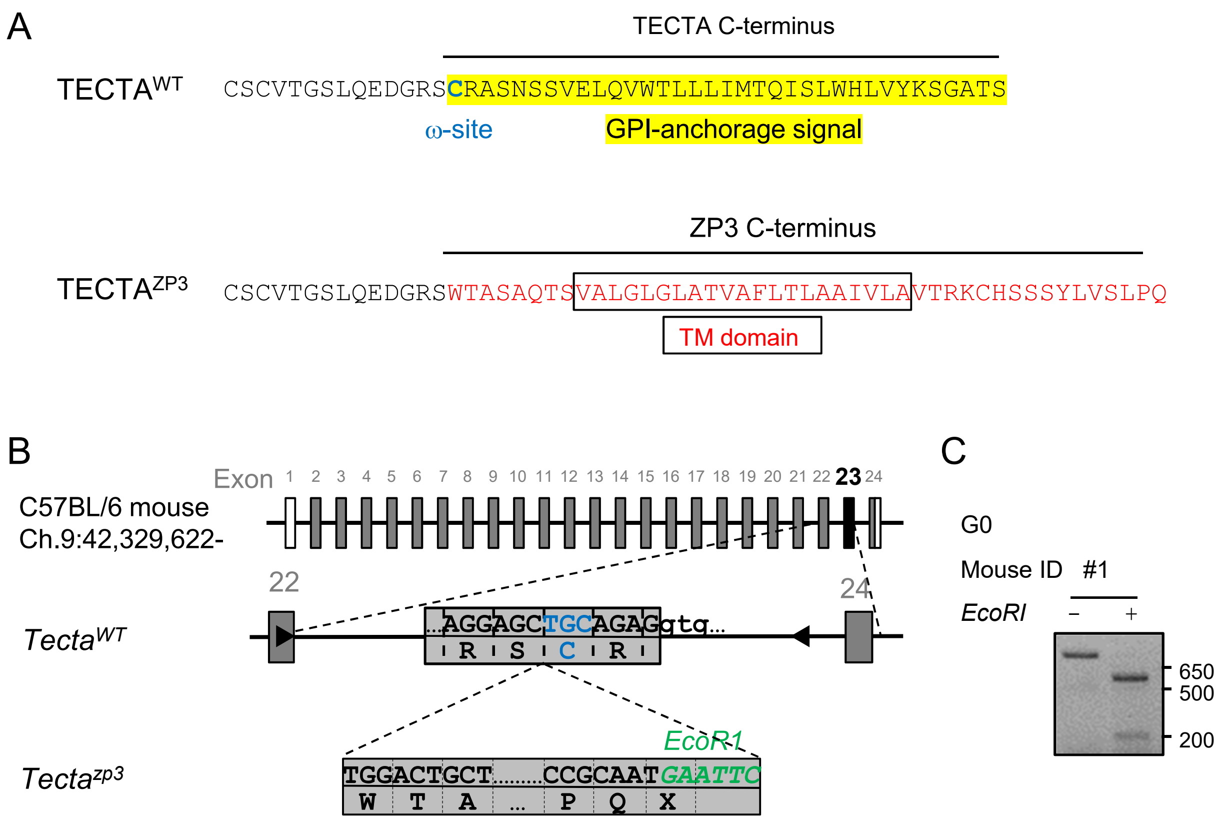
A
TECTA C-terminus
TECTAWT CSCVTGSLQEDGRSCRASNSSVELQVWTLLLIMTQISLWHLVYKSGATS
ω-site
GPI-anchorage signal
ZP3 C-terminus
TECTAZP3 CSCVTGSLQEDGRSWTASAQTSVALGLGLATVAFLTLAAIVLAVTRKCHSSSYLVSLPQ
TM domain
B
Exon 1 2 3 4 5 6 7 8 9 10 11 12 13 14 15 16 17 18 19 20 21 22 23 24
C57BL/6 mouse
Ch.9:42,329,622-
22
24
TectaWT
...AGGAGCTGCAGAGgtg...
R S C R
Tectazp3
EcoR1
TGGACTGCT.........CCGCAATGAATTC
W T A ... P Q X
C
G0
Mouse ID #1
EcoRI – +
650
500
200

**Figure S6. Related to Figure 6**

A. C-terminal protein sequence of TECTA and ZP3. The GPI-anchorage sequence including the omega site (blue) is replaced with the C-terminus of ZP3, including the transmembrane (TM) domain and a cytoplasmic tail.

B. Generation of *Tecta*$^{ZP3/ZP3}$ mice by CRISPR/Cas-9. The omega site (blue) is replaced with a donor sequence containing ZP3 C-terminus, stop codon, and EcoRI site.

C. PCR followed by EcoRI treatment showed the targeted mutation in the G0 founders.
